# A Bear-Specific Coding Exon in TMEM41B Is Associated with Cold Adaptation

**DOI:** 10.64898/2026.01.13.699376

**Authors:** Shuang Zhang, Hui Wang, Xinyi Zhang, Xiaodan Lu, Yu Xu, Kexin Xu, Jiaqi Zhang, Yuxuan Chen, Yonghao Feng, Yuru Wang, Yan Liu, Penglai Zou, Hanyin Zhang, Jiajun Yao, Yue Ma, Yanchun Xu, Hongying Ye, Shaohua Fan, Yinqi Bai, Tiemin Liu, Xingxing Kong

## Abstract

Bears exhibit an unusual form of hibernation in which core body temperature remains relatively high despite profound suppression of whole-body metabolism. How such a warm yet hypometabolic state is achieved remains poorly understood. Here, through integrative analysis of 238 RNA-seq datasets spanning seven species, we identify transmembrane protein 41B (TMEM41B) as an evolutionarily conserved factor associated with thermogenic capacity. Loss of Tmem41b in mice and Drosophila results in severe cold intolerance, whereas its overexpression enhances thermogenic performance, supporting a conserved role in heat production. Comparative genomic analyses reveal a bear-specific 51_–_amino acid N-terminal extension in TMEM41B, arising from a 5_′_ untranslated region mutation that converts a premature stop codon into coding sequence. Interactome profiling indicates that this bear-specific isoform differs from the canonical TMEM41B in its protein interaction landscape, including reduced association with components of mitochondria-associated endoplasmic reticulum membranes such as voltage-dependent anion-selective channel protein 1 (VDAC1). Consistent with this altered interaction profile, the extended isoform is associated with reduced mitochondrial oxidative activity. Functional analyses further suggest that the 51_–_ amino acid extension modulates multiple cellular programs, including suppression of oxidative phosphorylation and myogenic differentiation alongside activation of osteogenic pathways, collectively biasing cells toward a hypometabolic state. Notably, expression of the polar bear TMEM41B isoform in mice induces a torpor-like phenotype under fasting conditions. Together, these findings identify TMEM41B as a conserved regulator of thermogenic capacity and suggest that evolutionary modification of its N terminus may contribute to metabolic suppression and energy conservation during bear hibernation.

## Introduction

Large hibernating mammals, such as bears, exhibit remarkable physiological adaptations that enable them to survive prolonged periods of fasting and extreme cold. During hibernation, bears drastically suppress metabolic rate, reduce energy expenditure, and shift fuel utilization toward lipid oxidation while preserving muscle mass and organ function (1). Unlike small hibernators that rely heavily on UCP1-dependent brown fat thermogenesis (*2, 3*), the bears maintain relatively high and stable body temperatures, relying on finely tuned mechanisms of metabolic suppression rather than profound hypothermia (*4, 5*). These capabilities imply the existence of specialized regulatory systems that balance energy conservation with cellular homeostasis, yet the molecular mechanisms enabling this balance remain largely unknown.

Thermogenesis is a defining physiological strategy that supports survival, ecological expansion, and adaptive evolution (*6*). By enabling organisms to counter environmental cold and maintain stable internal temperatures, thermogenic programs have shaped key transitions in vertebrate history. Much of what is known about these programs derives from mammalian and avian studies, which have established that both shivering and non-shivering thermogenesis (NST) are mobilized during cold exposure(*7–9*). Evolutionary genetic analyses have revealed that NST is underpinned by both the emergence of lineage-specific factors and the conservation of core metabolic regulators. Comparative genomics has revealed a mosaic of lineage-specific innovations, such as UCP1, alongside universally conserved metabolic regulators like PGC1α (*6, 10–12*). Despite decades of study, it remains unclear whether a unified regulatory logic governs thermogenesis across species, or whether specialized metabolic strategies—exemplified by hibernation—are enabled by previously unrecognized molecular adaptations.

TMEM41B, an endoplasmic reticulum (ER)-resident membrane protein, is known for its roles in autophagy (*13*), lipid scrabbling (*14*) and viral infections (*15*). Although these functions suggest a capacity to organize membrane structure, TMEM41B has never been linked to thermogenesis. Here, we combine cross-species transcriptomic analysis with functional genomics to identify TMEM41B is essential for cold resistance across distant species: loss of *Tmem41b* in mice or its homolog in *drosophila* leads to profound cold intolerance, whereas increasing its expression enhances thermal resilience. Notably, bears harbor a N-terminal coding extension of 51 amino acids (51aa), absent from primates and rodents. We show that this extension is expressed and functional, and that introducing a chimera of polar bear 51aa with human TMEM41B into mice mimicked a torpor-like performance during fasting. This evolutionary novelty, coupled with the conservation of the core protein, positions TMEM41B as a previously unrecognized regulator potentially contributing to lineage-specific thermogenic adaptations such as hibernation.

## Results

### Phylogenetic analysis of non-shivering thermogenesis

Independent acquisitions of thermogenic strategies have long been central to palaeophysiological hypotheses regarding the origin of and diversification of species (*16*). After collecting and classifying these strategies from public literature (*6, 9, 17–19*) within a phylogenetic framework, we reviewed the mechanisms of NST as they progressively emerged in various vertebrate tissues under evolutionary constraints (Fig. 1A). To link NST to its underlying cellular and molecular mechanisms, we curated 238 high-quality bulk RNA datasets from 7 species comprising treatment-versus-control pairs reported in the public literature (Fig. 1B, table S1). From these datasets, we identified the top 5% of the most frequently occurring differentially expressed genes (DEGs) (table S2).

**Fig. 1.**
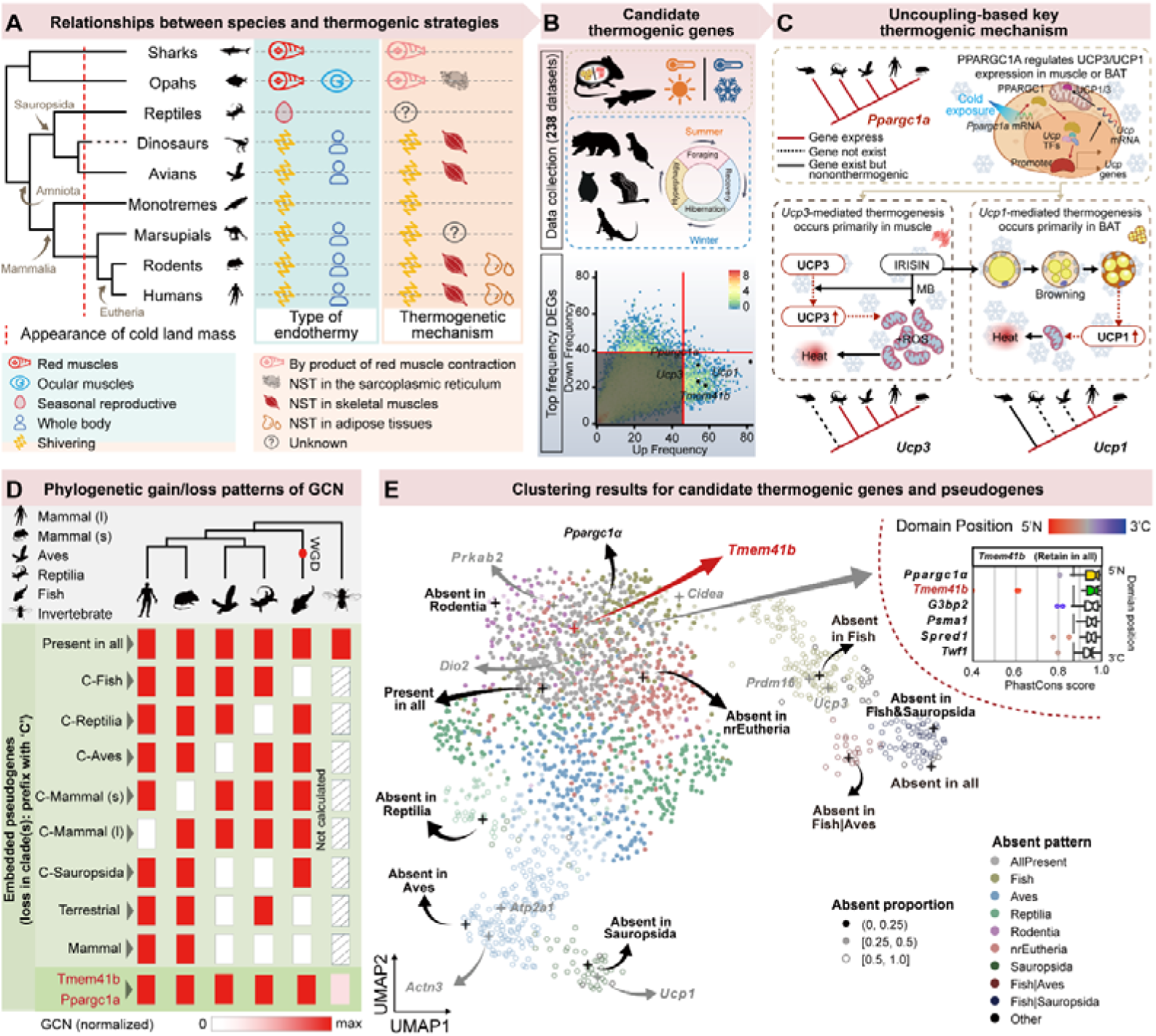
Phylogenetic study of candidate thermogenic genes. (**A**) Summary of evolutionary relationships in species phylogeny and physiological strategies of thermogenesis. Information on physiological strategies and mechanisms is integrated from public literature. The red dashed line indicates the emergence of cold land masses in the Early Cretaceous, after which endothermic mechanisms began to confer a selective advantage. NST, non-shivering thermogenesis. (**B**) Schematic diagram illustrating the curation of candidate thermogenic genes that are differentially expressed under cold-exposure or active-hibernation conditions. Top DEGs are compiled from 41 related research projects and 238 paired datasets. Known genes (*Ppargc1*α, *Ucp1*, and *Ucp3*) involved in the uncoupling of oxidative phosphorylation are highlighted. (**C**) Schematic diagram illustrating the molecular orchestration of *UCP* genes for thermogenesis and the complementary patterns observed in the corresponding gene gain/loss trees. (**D**) Phylogenetic gain and loss patterns of gene copy number (GCN) in representative thermogenic genes and pseudogenes. Only pseudogenes that are clustered with *bona fide* candidate thermogenic genes are retained. GCN is normalized within each clade, and there is a positive correlation between the intensity of red coloration and copy number. According to the minimum-evolution principle, a gain is defined as the continuous emergence of gene copies along the phylogenetic tree; all other patterns are considered losses in specific clades. Mammal (l): large mammals; Mammal (s): small mammals, most of which are rodents. (**E**) UMAP visualization of the clustering results for candidate thermogenic genes and pseudogenes. Pseudogenes are named according to their clade-specific gain/loss patterns. For instance, “Absent in *Aves*” indicates that the gene is exclusively lost in Aves while retained in all other clades, and “Present in all” indicates retention in all clades. Top right inset: Notched box plots displaying the top conserved genes in *Ppargc1*α (“Present in all”) clusters. The coding sequence of each gene is divided into 45-nucleotide-long domains, and the average phastCons conservation score is calculated using the median value. PhastCons scores represent the degree of evolutionary conservation at each coding domain, such that higher scores reflect stronger selective constraint across species. Dots are colored according to their domain positions from the 5’ end (red) to the 3’ end (blue). Outliers are additionally marked based on their domain loci within the gene. The dark-grey dashed line indicates the median conservation score in the gene cluster. Genes shown in grey represent known thermogenic gene *Ppargc1*α, while the gene highlighted in red, *Tmem41b* is the focus of this study.

We first verified that well-known genes involved primarily in the uncoupling of oxidative phosphorylation (i.e., via mitochondrial proton leak)—including peroxisome proliferator-activated receptor gamma coactivator 1-alpha (*Ppargc1*α), *Ucp1*, and uncoupling protein 3 (*Ucp3*)—were among the top DEGs (Fig. 1B). The molecular orchestration of these genes is highly interpretable through their complementary patterns observed in the corresponding gene gain/loss trees (Fig. 1C).

Therefore, to systematically explore additional genes functionally related to NST in vertebrates from an evolutionary and anthropology perspective, we hypothesized that candidate genes would: (i) undergo evolutionary changes that associated with diverse thermogenic mechanisms, as evidenced by gene gain/loss patterns traced throughout vertebrate evolution; and (ii) maintain high sequence conservation, thereby contributing to tissue specificity within their respective thermogenic clades. Notably, these criteria may be relaxed in teleosts, where lineage-specific whole-genome duplication (WGD) events have been documented (Fig. 1D) (*20*).

We then clustered all candidate genes based on a customized gene gain/loss metric (see Methods; Fig. 1E). Intriguingly, when we included phylogenetically constructed pseudogenes—encompassing all possible clade-wise gain/loss patterns—in the clustering analysis, ten of these pseudogenes clustered alongside *bona fide* gene neighborhoods in the Uniform Manifold Approximation and Projection (UMAP) embeddings (*21*). The result indicated that most real-world NST mechanisms can be cataloged into ten distinct phylogenetic frameworks. These clusters revealed that, in addition to being entirely lost or retained across all clades, candidate thermogenic genes may experience gene copy losses in specific clades (e.g., *fish*, *reptiles*, *aves*, small mammals, large mammals, or *Sauropsida*) or gains in non-avian terrestrial animals or mammals. Within these clusters, some are well-represented by the aforementioned uncoupling protein genes (*Ppargc1*α and *Ucp1)*. Others, however, are less studied, so we named these clusters based on their gain/loss patterns.

Subsequently, we calculated the conservation scores (*22*) of gene sequences in each clade (Fig. 1E, S1A). Although extensive studies have explored the evolution of thermogenic molecules such as UCP1 (*23*), these factors are present only in certain species. To date, little is known about whether there exist universally conserved genes that regulate thermogenesis across different taxa. The cluster corresponding to thermogenic transcription coactivator *Ppargc1*α is conserved during species evolution. Thus, this cluster was prioritized for analysis. Unsurprisingly, the *Ppargc1*α was the most conserved gene in the “Present in All” cluster. Interestingly, the second most conserved gene—the ER transmembrane protein TMEM41B—attracted our attention because its N-terminal region appears to have undergone relaxed selective pressure (Fig. 1E). This gene forms a conserved single-copy family across terrestrial vertebrates (6 out of 176 species in the Ensembl database) but exhibits relatively relaxed selection in *fish* (Fig. S1B). Moreover, its sequence underwent rapid evolution during the transition from aquatic to terrestrial vertebrates (Fig. S1C), which may reflect adaptation to changes in habitat. Previous studies indicated that TMEM41B plays roles in virus defense and energy metabolism (*24*). We therefore examined its effects on thermogenesis in a particular cellular context.

### Mice with *Tmem41b* knockout in adipocytes show impaired thermogenic response and mitochondrial function

Brown adiposse tissue (BAT), beige adipose tissue, and skeletal muscle are the three primary thermogenic tissues (*25*). These tissues play distinct yet interconnected roles in thermogenesis, contributing to energy homeostasis and metabolic adaptation. Studies have clearly identified adipose tissue as a key site for NST in mammals (*26*). To investigate whether TMEM41B influences thermogenesis, we firstly generated adipose tissue-specific *Tmem41b* knockout (*Tmem41b^AdiCre^*) mice by crossing the *adiponectin*-*Cre* mice with *Tmem41b* flox (*Tmem41b^f/f^)* mice (Fig. 2A, S2A, B). The growth of *Tmem41b^AdiCre^* mice, both male and female, was comparable to that of the *Tmem41b^f/f^*controls when maintained on a chow diet at room temperature (Fig. S2C, D). However, under cold stress (4°C for 4 hours), the *Tmem41b^AdiCre^* mice displayed a significant drop in core temperature, reaching approximately 20°C (Fig. 2B, movie 1). Similarly, the postnatal day 7 (D7) *Tmem41b^AdiCre^* pups were more susceptible to cold-induced hypothermia after 1-hour exposure (Fig. 2C). The biology of adipose tissue was then assessed. While the weight of white adipose tissue (WAT) showed no genetic differences, the BAT weight of *Tmem41b^AdiCre^* mice was significantly increased compared to controls (Fig. S2E). Histological analysis revealed larger lipid droplets in the BAT of *Tmem41b^AdiCre^* mice (Fig. S2F), with no noticeable changes in inguinal WAT (iWAT) or epididymal WAT (eWAT) (Fig. S2G). The transmission electronic microscope (TEM) further showed enlarged mitochondria with disrupted cristae in BAT (Fig. S2H). Functionally, BAT from *Tmem41b^AdiCre^* mice exhibited a reduced oxygen consumption rate (OCR) and diminished protein levels of mitochondrial complex subunits NDUFB8 and SDHB (Fig. S2I-K). Correspondingly, thermogenic gene expression was suppressed in the BAT of *Tmem41b^AdiCre^* mice (Fig. S2L), with reduced protein levels of UCP1 (Fig. S2M), a key protein for NST. In addition, bulk RNA sequencing (RNA-seq) analysis of BAT corroborated these findings, showing significant downregulation of thermogenic and respiratory pathways, but upregulation of phagosomes (Fig. S2N to P). Given that the signaling of phagosome was increased, we examined autophagy-related markers. Indeed, the protein levels of autophagy, including LC3, P62, PARKIN, were increased in BAT from *Tmem41b^AdiCre^* mice (Fig. S2Q). This is consistent with the notion that excessive activation of autophagy may impair the function of thermogenic cells and lead to a reduction in energy metabolism (*27*).

**Fig. 2.**
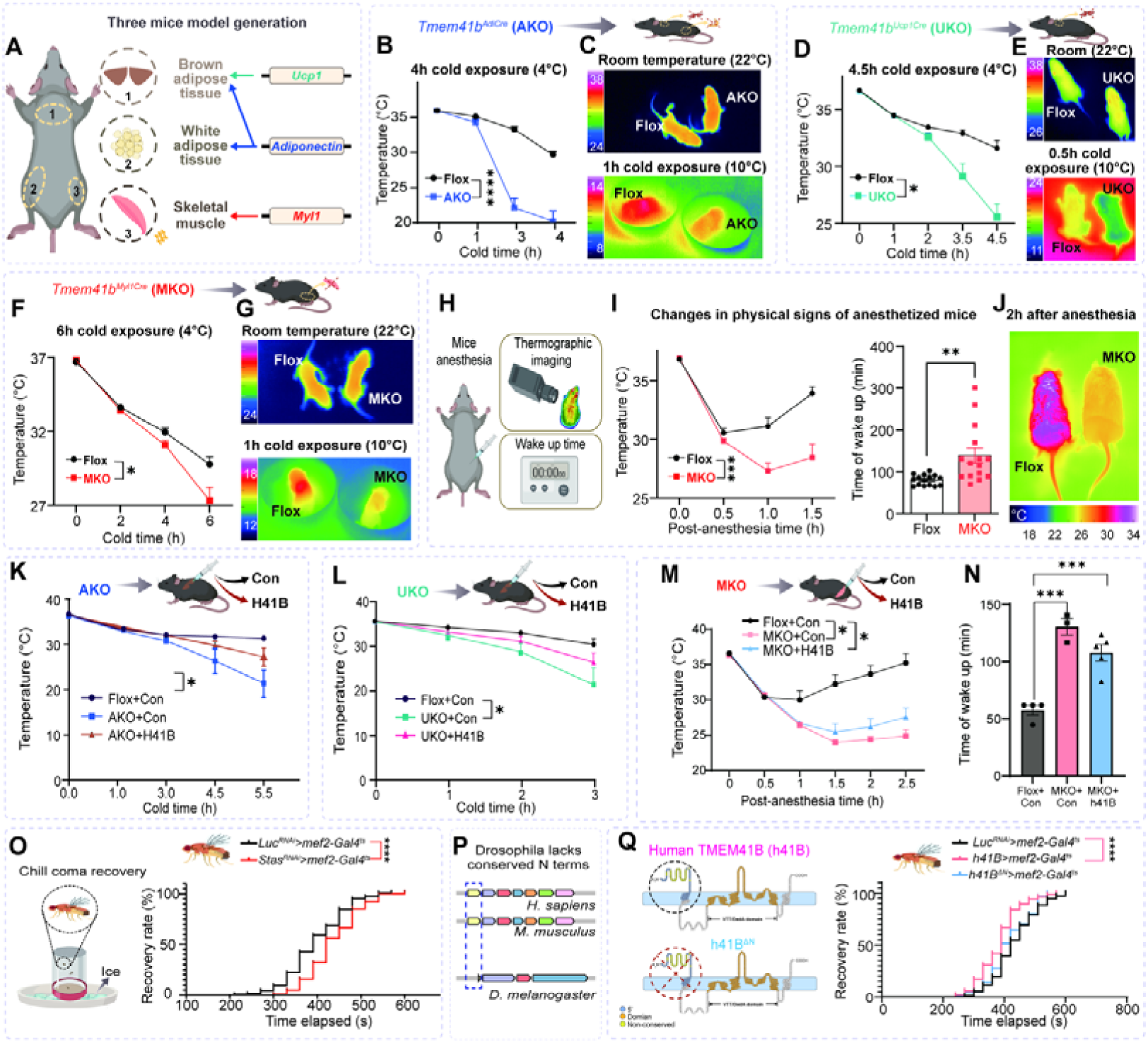
TMEM41B is a linchpin of thermoregulation across tissues and species. **(A)** Schematic illustration depicting the three primary thermogenic tissues in mice. (**B**) Body temperature curves of *Tmem41b^f/f^* (Flox) and *Tmem41b^AdiCre^* (AKO) mice during 4 hours of cold exposure (4°C). Data represent mean ± SEM (n = 7, 6 per group). (**C**) Infrared thermography images of 7-day old *Tmem41b^f/f^* and *Tmem41b^AdiCre^* mice at room temperature (RT, 22°C) and after 1 hour of cold exposure (10°C). Warmer regions appear in red/yellow, while cooler regions appear in blue. (**D**) Body temperature curves of *Tmem41b^f/f^* (Flox) and *Tmem41b^Ucp1Cre^* (UKO) mice during 4.5 hours of cold exposure (4°C). Data represent mean ± SEM (n = 9, 8 per group). (**E**) Infrared thermography images of 7-day old *Tmem41b^f/f^*and *Tmem41b^Ucp1Cre^* mice at RT (22°C) and after 0.5 hour of cold exposure (10°C). (**F**) Body temperature curves of *Tmem41b^f/f^* and *Tmem41b^Myl1Cre^* mice during 6 hours of cold exposure (4°C). Data represent mean ± SEM (n = 11 per group). (**G**) Infrared thermography images of 7-day old *Tmem41b^f/f^*and *Tmem41b^Myl1Cre^* mice at RT (22°C) and after 1 hour of cold exposure (10°C). (**H-I**) The temperature of indicated mice after anesthesia (n = 15 per group) (left). The wake-up time of mice (n = 15 per group) (right). (**J**) Infrared thermographic imaging of mice 2 hours after anesthesia. (**K-L**) Cold tolerance test of BAT specific reconstitution of human TMEM41B in Flox, AKO and UKO mice (n = 3 - 4 per group). (**M-N**) Body temperature curves and the wake-up time of muscle-specific reconstitution of human TMEM41B in Flox and MKO mice (n= 3 - 5 per group). (**O**) *Drosophila* were subjected to 20 minutes of cold exposure at 0℃ and monitored for recovery (n > 100, log-rank test). *Stasimon* knockdown *Drosophila* exhibited prolonged recovery times from acute cold exposure compared to controls. (**P**) Schematic representation of the protein domain architecture of TMEM41B orthologs in human, mice, and *Drosophila* melanogaster, highlighting the absence of the N-terminal region (N) in the insect ortholog. (**Q**) Expression of full-length human TMEM41B (*UAS-hTMEM41B*) *Drosophila* rescued cold stress recovery, whereas the C-terminal domain alone (*UAS-hTMEM41B*^Δ*N*^) failed to restore normal recovery times. Residues within the known VTT/DedA domain are marked with orange circles, and residues are highlighted in yellow.

To exclude potential contributions of heat production from WAT, we generated brown adipocyte-specific *Tmem41b* knockout (*Tmem41b^Ucp1Cre^*) mice (Fig. 2A, S3A, B). Both male and female *Tmem41b^Ucp1Cre^* mice exhibited growth patterns comparable to *Tmem41b^f/f^* controls on a chow diet at room temperature (Fig. S3C, D). In contrast, under cold stress (4°C for 4.5 hours), *Tmem41b^Ucp1Cre^* mice experienced a marked drop in core temperature, reaching 25 °C for male and 20 °C for female (Fig. 2D, S3E, movie 2). Furthermore, D7 *Tmem41b^Ucp1Cre^*pups were more susceptible to cold-induced hypothermia after just 0.5 hour of exposure (Fig. 2E). Consistent with the observations in *Tmem41b^AdiCre^* mice, *Tmem41b^Ucp1Cre^* mice exhibited no significant differences in WAT weight but showed a significant increase in BAT weight (Fig. S3F). Histological analysis revealed larger lipid droplets in BAT, whereas the changes in iWAT and eWAT did not reach statistical significance (Fig. S3G, H). Moreover, mitochondrial morphology in BAT was compartmentalized, as observed using TEM (Fig. S3I). BAT from *Tmem41b^Ucp1Cre^* mice exhibited reduced OCR and decreased levels of mitochondrial complex subunits, including NDUFB8 and SDHB (Fig. S3J, K). Thermogenic gene expression in BAT was also significantly downregulated (Fig. S3L), accompanied by lower UCP1 protein levels (Fig. S3M). Collectively, these findings highlight the critical role of TMEM41B in regulating BAT structure and function, particularly in thermogenesis and adaptation to cold exposure (Fig. S3N).

### Muscle-specific loss of *Tmem41b* produces cold intolerance in both mice and drosophila

Notably, skeletal muscle NST appears to be the more ancestral form and is more closely associated with large mammals, such as adult humans (*9*). We next generated *Tmem41b^Myl1Cre^* mice (Fig. 2A, S4A-C) to investigate the thermogenic role of TMEM41B in muscle. *Tmem41b^Myl1Cre^*mice exhibited normal growth rates comparable to their *Tmem41b^f/f^*littermates up to 10 weeks of age on a standard chow diet at room temperature (Fig. S4D-F). Moreover, these mice showed similar energy expenditure between genetics (Fig. S4G-J). However, the *Tmem41b^Myl1Cre^*mice were cold sensitive, with their core body temperature dropping to 27°C within 6 hours of cold exposure (Fig. 2F). While the skin temperature of 7-day old pups was comparable among littermates at room temperature, the *Tmem41b^Myl1Cre^* pups experienced exacerbated cold stress-induced hypothermia (Fig. 2G). Both shivering and non-shivering mechanisms are known to contribute to muscle-driven thermogenesis during cold exposure (*28*). Thus, we performed an electromyography test, the gold standard for shivering assessments (*28*), to evaluate muscle shivering and found no significant differences between the genetic models (Fig. S5A). To exclude contributions from shivering-induced thermogenesis, mice were anesthetized prior to testing. Under these conditions, the core temperature of *Tmem41b^Myl1Cre^* mice dropped faster but recovered more slowly than *Tmem41b^f/f^* mice (Fig. 2H, I). The infrared thermographic imaging further confirmed hypothermia in *Tmem41b^Myl1Cre^* mice 2 hours post anesthesia (Fig. 2J). While impaired thermogenesis in skeletal muscle likely underlies the cold sensitivity of *Tmem41b^Myl1Cre^* mice, other thermogenic tissues, such as BAT and beige adipose tissue—key players in NST (*29*)—might also contribute to this phenotype. To investigate this further, we assessed the thermogenic capacity of adipose tissue. Histological analysis of BAT showed no significant differences between genotypes. However, the iWAT from *Tmem41b^Myl1Cre^*mice surprisingly exhibited an increased number of multilocular cells (Fig. S5B). This finding was supported by elevated UCP1 protein levels (Fig. S5C), suggesting a compensatory mechanism in iWAT to offset their impaired heat production in muscle. These data suggested that the cold sensitivity of *Tmem41b^Myl1Cre^* mice stemmed from skeletal muscle dysfunction rather than dysregulation of adipose tissue NST.

To elucidate the pathways altered in skeletal muscle from *Tmem41b^Myl1Cre^* mice, we performed bulk RNA-seq. There were 3917 different expression genes in *Tmem41b^Myl1Cre^* mice compared to *Tmem41b^f/f^* mice (Fig. S5D, E). Among the DEGs, 2,335 genes were up-regulated and 1,582 genes were down-regulated in *Tmem41b^Myl1Cre^* mice (Fig. S5E). Kyoto Encyclopedia of Genes and Genomes (KEGG) enrichment revealed a significant enrichment of the pathways associated with inflammatory pathways and a decrease in cellular respiration (Fig. S5F, G), consistent with transcriptomic changes observed in BAT. RT–qPCR further validated the upregulation of inflammatory genes upon *Tmem41b* deletion (Fig. S5H).

Given that mitochondria serve as central hubs for heat production (*30*), we next examined whether the thermogenic defects caused by TMEM41B deficiency arise from impaired mitochondrial structure or function. Transmission electron microscopy showed that *Tmem41b^Myl1Cre^*myofibers contahaodined mitochondria that were swollen and exhibited severely disrupted cristae (yellow arrows, Fig. S6A), although mitochondrial abundance remained unchanged (Fig. S6B). Notably, a subset of mitochondria displayed onion-like concentric membrane structures, a hallmark of severe ultrastructural abnormalities (Fig. S6C). Concordantly, the protein levels of SDHB and NDUFB8 were markedly reduced (Fig. S6D), accompanied by a significant decrease in OCR in *Tmem41b*-deficient muscle (Fig. S6E, F). In line with this, the primary myoblasts from *Tmem41b^Myl1Cre^* mice displayed fragmented, in contrast to the highly interconnected and elongated mitochondrial network observed in controls (Fig. S6G). Further analysis revealed that the mitochondrial membrane potential was diminished in *Tmem41b*-deficient myocytes (Fig. S6H). To confirm these findings in a cell-autonomous way, we employed the CRISPR/Cas9 knockout system to delete *Tmem41b* (single guide *Tmem41b*, sg41B) in C_2_C_12_ myoblast cell line (Fig. S6I). The CRISPR knockout cells exhibited reduced mitochondrial protein levels and attenuated OCR, consistent with observations in primary myocytes (Fig. S6J-L). Taken together, these results indicated that the loss of TMEM41B disrupted mitochondrial structure and function, impairing OXPHOS activity and membrane potential.

To further elucidate the biological significance of the TMEM41B in cold tolerance, we performed bilateral injections of human TMEM41B (*AAV-hTmem41b*) into the interscapular BAT of *Tmem41b^f/f^*, *Tmem41b^AdiCre^* (Fig. 2K, S7A) and *Tmem41b^Ucp1Cre^*(Fig. 2L, S7B) mice and GAS muscle of *Tmem41b^f/f^* and *Tmem41b^Myl1Cre^* mice (Fig. 2M, N, S7C). Notably, overexpression of TMEM41B successfully rescued the cold intolerance phenotypes in AKO, UKO and MKO mice models, mitochondrial respiration and restored the expression levels of UCP1, SDHB, and NDUFB8 proteins (Fig. S7D-F), underscoring the functional importance of TMEM41B in thermogenic adaptation.

To explore the evolutionary relevance of TMEM41B-mediated thermogenesis, we used *mef2-Gal4^ts^* driver line to generate the *stasimon* (*Drosophila* homolog of *Tmem41b*) knockdown fruit flies and evaluated their response to cold stress using the chill coma recovery assay (*31*) (Fig. 2O). *Stasimon* knockdown flies exhibited a significantly prolonged chill coma recovery time compared to controls, indicating impaired cold tolerance (Fig. S8A). Although TMEM41B exhibits a conserved role in thermogenesis, its protein structure differs between humans and *Drosophila*, particularly in the length of the first N-terminal transmembrane (TM) segment, which is longer in humans (Fig. 2P). To determine whether this first TM domain contributes to thermogenic function, we generated constructs expressing either full-length human TMEM41B (hTMEM41B) or an N-terminal–truncated variant (hTMEM41B^ΔN^) (Fig. 2Q, S8B). In the chill-coma recovery assay, only full-length human TMEM41B enhanced thermogenic capacity in flies, whereas the N-terminal–truncated form showed no effect (Fig. 2Q, S8C). These results indicate that the first N-terminal transmembrane domain is sufficient, and the conserved C-terminal domains are essential, for thermogenesis.

### Bear TMEM41B contains an additional 51 amino acids at its N-terminus

The C-terminus of TMEM41B is highly conserved across species while the N-terminus is variable (Fig. 1E). This pattern suggests that the conserved C-terminal region encodes core ancestral functions, while diversification of the N-terminus may have provided an evolutionary substrate for lineage-specific metabolic adaptations. Next, we conducted analyses on the conservation of TMEM41B proteins. Consistent with this idea, our comparative analysis revealed that, in contrast to the highly conserved VTT domain (also referred to as the DedA domain) at the C-terminus—which forms a superfamily with bacterial DedA family proteins sharing evolutionary origin (*32*)—the N-terminal residues upstream of the first transmembrane loop exhibit remarkable plasticity (Fig. S9A, B).

Within this variable region, we identified a previously unannotated coding exon located upstream of the canonical TMEM41B start site that is present in the *Ursidae genus* (Fig. 3A, Fig. S10A-B, table S3). Notably, a T-to-C substitution at the 49th nucleotide converts the ancestral stop codon (TGA) into an arginine codon (CGA) in multiple *Ursidae* species (including *Ursus thibetanus*, *U. maritimus*, *U. arctos*, *U. americanus*, *Helarctos malayanus*, and *Ailuropoda melanoleuca*), with the exception of *Tremarctos omatus*, which carries a T-to-G mutation that converts the stop codon into a glycine (GGA) (Fig. 3B, Fig. S10C). The consistent loss of the ancestral stop codon across nearly all extant bear species indicates that this enabling mutation likely arose in the common ancestor of bears between ∼40 and 19.26 Mya (Fig. 3B). Importantly, the canonical exon–intron boundary between this newly identified upstream exon and the downstream coding region is preserved, supporting its incorporation into the mature transcript. Furthermore, RT-qPCR and transcriptomic data across 4 bear species indicate that this novel exon is transcriptionally active (Fig. 3C, D, table S4). To verify bears produce a longer TMEM41B isoform than other vertebrates, we compared RNA and protein from bears with those of related mammals, including foxes. Although both species showed detectable 5′ Tmem41b transcripts (Fig. 3D), only bears produced a TMEM41B protein containing the extended N-terminus (Fig. 3E), confirming that bears uniquely express the long TMEM41B isoform.

**Fig. 3.**
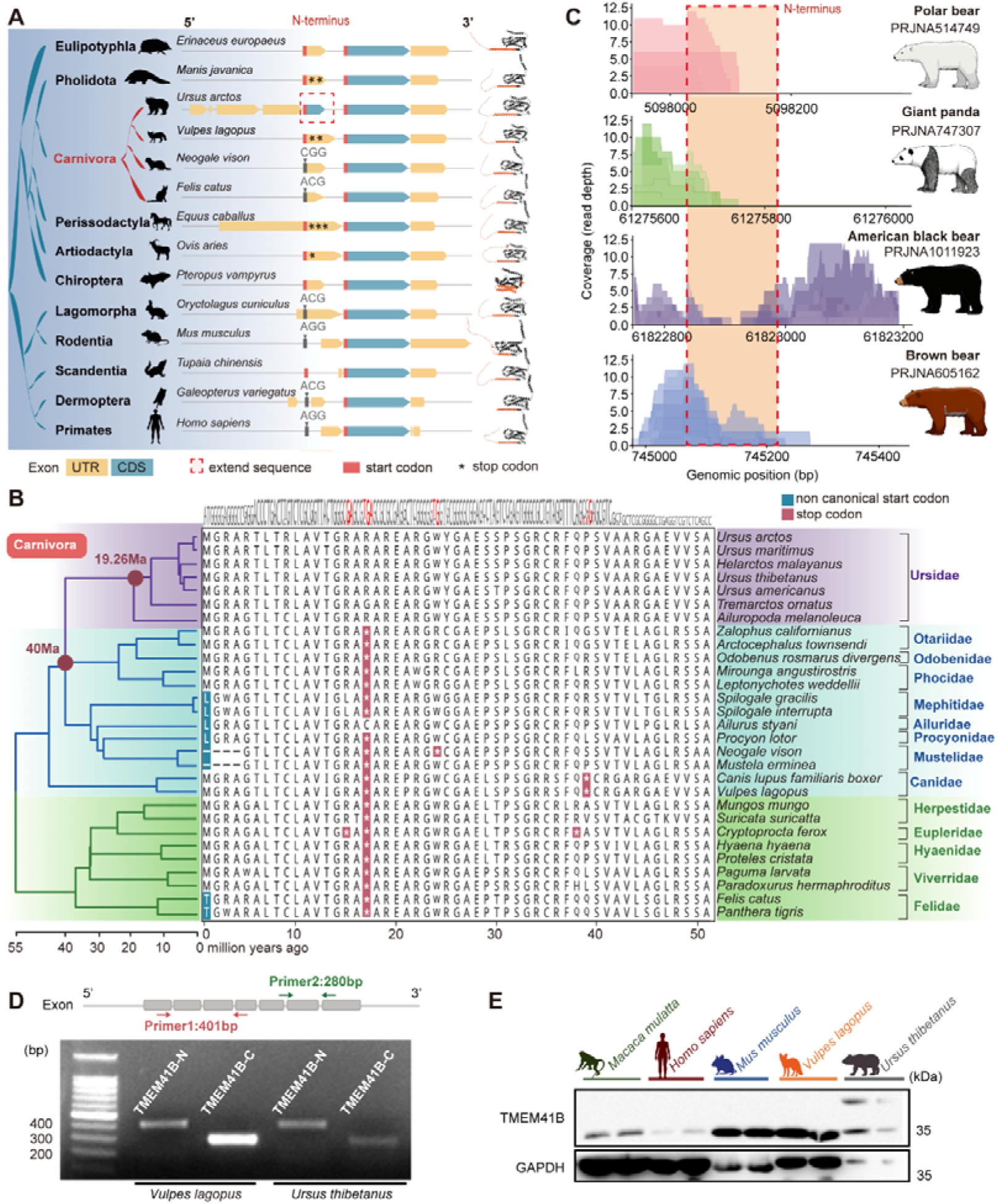
Evolutionary diversification of the TMEM41B N-terminal region across Boreoeutheria. (**A**) Comparative gene and protein structures of TMEM41B across representative Boreoeutheria lineages. Each row represents one species, showing the exon–UTR–CDS organization and the corresponding protein schematic. A pronounced N-terminal coding region extension is observed specifically in Ursidae (red dashed box). Non-canonical start codons and premature stop codons are indicated. See Fig. S10A, table S3 for complete species lists and underlying genomic sequences. (**B**) Multiple sequence alignment of the TMEM41B N-terminal amino acid sequences from 30 carnivoran species. The phylogenetic tree and divergence times were obtained from TimeTree (https://timetree.org/). Non-canonical start codons (blue) and lineage-specific premature stop codons (red) are highlighted. Ursidae exhibits a restored open reading frame due to two fixed substitutions in this region, in contrast to the truncated N-terminal sequences observed in most other carnivorans. (**C**) RNA-seq read coverage across the *TMEM41B* N-terminal region in four bear species. Each color represents an independent RNA-seq sample, with corresponding sample accessions shown in the legend. The shaded pink region indicates the N-terminal segment analyzed in this study. (**D**) PCR confirmation of N and C terminal of *Tmem41b* in *Vulpes lagopus* and *Ursus thibetanus*. (**E**) Western blot analysis of TMEM41B expression in muscles from indicated species.

### 51-amino-acids substitution induces a torpor-like state in mice

To examine the functional significance of the additional N-terminal region in bears, we generated a chimeric construct (B41B) by inserting the extra 51aa from polar bear TMEM41B into the start site of the human TMEM41B (H41B) (Fig. 4A). Subcellular localization assays revealed that B41B predominantly resides in the endoplasmic reticulum (Fig. S11A). Using TurboID-based proximity labeling coupled with mass spectrometry in HEK293T cells, pathway analysis of the H41B and B41B interactomes revealed that H41B specifically interacts with mitochondria-associated ER membrane (MAM) proteins, whereas this interaction is almost lost in B41B (Fig. 4B, table S5). Co-immunoprecipitation (Co-IP) assays further confirmed that H41B, but not B41B, physically interacts with Voltage-dependent anion-selective channel protein 1 (VDAC1) (Fig. 4C), a member of MAM proteins. Concomitant with this, co-localization assays demonstrated that H41B is capable of co-localizing with mitochondria, whereas B41B is not (Fig. 4D). Considering that 41B is an ER-resident protein, these findings indicate that only H41B can localize to the MAM. These data suggest that the N-terminal extension may sterically hinder MAM association.

**Fig. 4.**
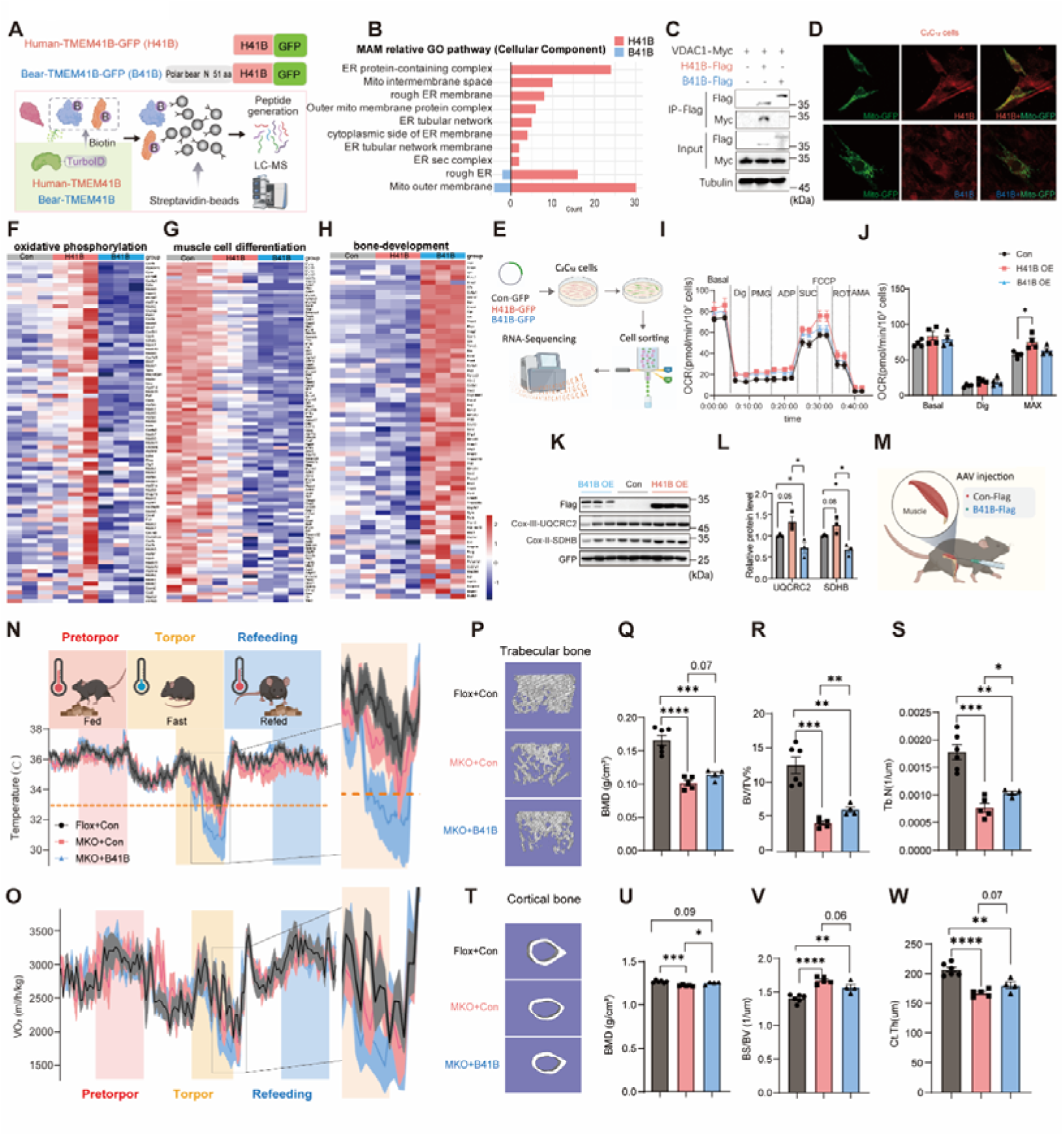
51-amino-acids substitution induces a torpor-like state in mice. (**A**) Turbo-ID-based proximity labeling combined with mass spectrometry to identify the interactome of Human TMEM41B (H41B) and Bear TMEM41B (B41B). (**B**) Biological Process terms from the Gene Ontology (GO) analysis for proteins identified in the TMEM41B-centric interactome related to MAM pathway. (**C**) Co-IP analysis confirms a specific protein-protein interaction between H41B and the mitochondrial outer membrane protein VDAC1, but not B41B and VDAC1. (**D**) Representative subcellular localization microscopy images of both H41B and B41B colocalizes with Mitochondria. Scale bar = 10 µm. (**E**) Schematic illustrating the generation of over-expression H41B or B41B C_2_C_12_ cells, fluorescence-activated cell sorting (FACS) and subsequent RNA sequencing (RNA-Seq) analysis. (**F-H**) Heatmap of RNA-seq. (F) Oxidative phosphorylation related genes; (G) Muscle cell differentiation related genes; (H) Bone-development related genes. (**I**) Representative traces of oxygen consumption rates (OCR) in over-expression H41B or B41B C_2_C_12_ cells (n = 4 per group). (**J**) Quantification of basal, C1, C2 and Maximum (n = 4 per group). (**K**) Western blot analysis of the mitochondrial complex SDHB and UQCRC2 in over-expression H41B or B41B C_2_C_12_ cells (n = 3 per group). (**L**) Corresponding densitometric quantification of protein levels. (**M**) Strategy for AAV-mediated delivery of bear TMEM41B into skeletal muscle of knockout mice). (**N-O**) The plot shows body temperature and VO_2_ in mice over 3 days while placed in TSE cages for a metabolic assay of processes pertinent to hibernation. The assay including pre-torpor phase (pink), torpor phase (organge, food deprived), and post-torpor ad libitum refeeding phase (blue). (N) Body temperature; (O) VO_2_. (**P-S**) Quantification of trabecular bone micro-architecture. (P) Micro-CT of trabecular bone; (Q) Bone Mineral Density (BMD);(R) Percentage of Bone Volume/Total Volume (BV/TV%); (S) Trabecular number (Tb.N). (T-W) Analysis of cortical bone micro-CT. (T) Micro-architecture of cortical bone; (U) BMD; (V) Bone Surface / Bone Volume (BS/BV); (W) Cortical Thickness (Ct.Th).

To determine the functional significance of the additional 51 amino acids, we overexpressed either B41B-GFP or human TMEM41B (H41B-GFP) in C_2_C_12_ cells. GFP-positive cells were isolated by flow cytometry and subjected to RNA-seq analysis (Fig. 4E). Principal component analysis (PCA) revealed distinct transcriptional profiles between B41B- and H41B-expressing C_2_C_12_ cells (Fig. S11B). Gene ontology (GO) analysis showed that B41B overexpression modestly downregulated oxidative phosphorylation and muscle cell differentiation, while upregulating pathways related to bone development compared with H41B (Fig. 4F-H, S11C, table S6). Consistently, OCR was elevated in H41B-overexpressing cells, whereas no significant change was detected in B41B-overexpressing cells relative to the control (Fig. 4I, J). The levels of several mitochondrial proteins were increased in H41B-expressing cells but reduced in B41B-expressing cells (Fig.4K, L). Collectively, these data suggest that the additional 51 amino acids in bear TMEM41B help maintain cells in a low metabolic state.

To further investigate the physiological function of the additional 51 amino acids, we administered AAV-B41B to mice to assess its *in vivo* effects. Because large adult mammals possess limited BAT and rely primarily on skeletal muscle for thermogenic regulation, intramuscular injection was performed in MKO mice, allowing direct evaluation of muscle-specific pathways (Fig. 4M, S12A-C). Remarkably, mice overexpressing B41B displayed an enhanced ability to maintain a hypothermic state under combined cold and fasting conditions (Fig. S12D). To further characterize the effect of B41B on torpor, we subjected mice to a 24-hour fasting period to mimic a hibernation-like state(*34, 35*). This treatment revealed that B41B-induced hypothermia was accompanied by a marked reduction in oxygen consumption rate (VO₂), and the decline in VO₂ closely paralleled the decrease in core temperature (Tcore) after fasting (Fig. 4N, O). Moreover, B41B-overexpressing mice exhibited a protective effect on bone (Fig. 4P-W, Fig. S12E-H). Together, these findings indicate that the additional 51 amino acids in bear TMEM41B promote a hypometabolic, torpor-like state *in vivo*.

## Discussion

This study identifies TMEM41B as an evolutionarily conserved regulator of thermogenic capacity and reveals how subtle sequence diversification can fundamentally reprogram metabolic strategies across species. The requirement of TMEM41B for cold resistance in both *Drosophila* and mice, together with the enhancement of thermogenic capacity upon its overexpression, demonstrates that TMEM41B functions as a deeply conserved component of adaptive thermogenesis—distinct from lineage-specific thermogenic factors such as UCP1. Protein sequence analysis further revealed that bears possess additional 51 amino acids at the N-terminus of TMEM41B, likely representing an adaptive modification linked to the evolution of hibernation. Consistent with this notion, overexpression of the polar bear TMEM41B isoform in mice induced hibernation-like phenotypes, including reduced body temperature and metabolic suppression during cold exposure.

Comparative analysis of *TMEM41B* across carnivoran lineages highlights substantial evolutionary plasticity in the N-terminal region, which contrasts with the otherwise high sequence conservation observed in the coding domain. The lineage-specific divergence observed in ursids suggests that the N-terminal segment may have been a hotspot for adaptive modifications, potentially contributing to functional specialization within the family. For example, the presence of a non-canonical ACG codon at the ancestral start site in *Felidae*, versus a canonical ATG in *Ursida*e, implies that translational efficiency and regulatory mechanisms at the N-terminus may have undergone distinct selective pressures across *carnivoran* lineages. In *ursids*, subsequent mutations restoring a continuous open reading frame likely refined N-terminal architecture, which could underlie lineage-specific functional divergence of *TMEM41B*.

The 51–amino acid N-terminal extension of the bear TMEM41B isoform profoundly alters its subcellular behavior and functional output. Mechanistically, this extension disrupts the protein’s ability to associate with MAM, including interactions with key components such as VDAC1. ER–mitochondria communication is attenuated, likely impairing calcium transfer and lipid flux, two critical processes that normally promote mitochondrial activity and energy expenditure (*36, 37*). By modulating ER-mitochondria tethering, TMEM41B effectively acts as a molecular “thermostat,” controlling the flow of signals and substrates that determine metabolic output. This provides a plausible explanation for the hypometabolic phenotype induced by the bear isoform at the cellular level.

From an evolutionary perspective, TMEM41B appears to remain indispensable for energy metabolism and thermogenesis. Intriguingly, hibernating species such as bears (*38*) harbor two TMEM41B isoforms of different lengths. Why these two forms coexist remains unclear. One plausible explanation is that the shorter isoform—which is similar in length to the human protein-supports thermogenic function during the active, non-hibernating season, whereas the bear-specific long isoform may act preferentially during hibernation to counterbalance the thermogenic activity of the short isoform and help maintain a hypometabolic state. How the two isoforms are regulated during evolution remains unclear; they may interact in ways analogous to other isoform pairs, such as CHREBP-1α and CHREBP-1c (*39*), where one isoform modulates the activity of the other. These evolutionary and mechanistic hypotheses will require further experimental validation.

Finally, current reference genome assemblies and gene annotations introduce uncertainty in interpreting N-terminal variation. The recent annotation of the N-terminal region as protein-coding in the brown bear, contrasted with its 5′ UTR designation in other ursids, likely reflects limitations of genome assembly and annotation rather than genuine biological differences. This underscores the importance of incorporating high-quality long-read genome assemblies and tissue-resolved transcriptomic data to accurately reconstruct gene architecture and evolutionary trajectories.

In summary, our study establishes TMEM41B as a conserved regulator of thermogenesis. The N-terminal extension of polar bear repurposes TMEM41B from promoting energy expenditure to facilitating metabolic suppression, providing a molecular basis for hibernation and energy conservation. These findings highlight how minor genetic changes in conserved proteins can drive profound shifts in organismal metabolism, linking molecular evolution to ecological adaptation, and offer potential avenues for therapeutic modulation of energy balance in metabolic disorders.

## Methods

### Animals

All animal experiments were conducted at Fudan University, following the guidelines set by the Science Research Ethics Committee (IDM2023034). *TMEM41B^flox/flox^* mice (Flox) were obtained from GemPharmatech Laboratory Animal Co., Ltd. (Jiangsu, China). **Animal Model 1**: Adipose Tissue-Specific TMEM41B Knockout. To investigate the role of TMEM41B in adipose tissue, we established adipose tissue-specific TMEM41B knockout mice by breeding *TMEM41B^flox/flox^* mice with *Adiponectin-Cre* transgenic mice. **Animal Model 2**: Brown Fat-Specific TMEM41B Knockout. To assess the function of TMEM41B in brown fat tissue, we generated brown fat-specific TMEM41B knockout mice by crossing *TMEM41B^flox/flox^* mice with *UCP1-Cre* transgenic mice. **Animal Model 3**: Skeletal Muscle-Specific TMEM41B Knockout. To examine the physiological role of TMEM41B in skeletal muscle, we generated skeletal muscle-specific TMEM41B knockout mice by crossing *TMEM41B^flox/flox^*mice with *Myl1-Cre* transgenic mice. **Animal Model 4 to 6**: To further investigate the functional of human TMEM41B in adipose tissue, brown adipose tissue (BAT) and skeletal muscle, adeno-associated viruses (AAVs) expressing human full-length TMEM41B (AAV-hTMEM41B) were packaged and purified. Viral particles were bilaterally injected into the interscapular BAT of 8-week-old *Tmem41b* AKO mice or *Tmem41b* UKO mice, and into the gastrocnemius (GAS) muscles of *Tmem41b* MKO mice. Control animals received AAV encoding GFP. Tissues were collected for histological and functional analyses following a 2-week recovery for *Tmem41b* AKO mice or *Tmem41b* UKO mice or a 4-week recovery for *Tmem41b* MKO mice. **Animal Model 7:** To evaluate the role of bear-TMEM41B in muscle tissue, the muscles of *TMEM41B^flox/flox^* and *TMEM41B^Myl1-cre^*mice were injected with an AAV vector expression a bear Tmem41b ortholog (B41B) or AAV control vector (Con). Functional and morphological rescue was assessed 2 weeks post-injection.

### Fly stains

Flies were reared in standard vials (25 mm × 95 mm) containing cornmeal medium and housed in a humidified, temperature-controlled incubator under a 12:12 light-dark cycle at 25L. UAS-*Luciferase^RNAi^* (*Luc^RNAi^*) and *mef2-Gal4^ts^*fly strains were kindly provided by Dr. Xinhua Lin (Fudan University) as previously described (*40*). UAS-stasimonRNAi(SatsRNAi) (II) (60128) was obtained from the Bloomington Drosophila Stock Center (BDSC). To generate the *UAS-hTMEM41B* and *UAS-hTMEM41B*^ΔN^ transgenic flies, the coding sequence of human TMEM41B (NM_015012.4, ENST00000528080.6) and the C-terminal TMEM41B construct containing a mutated amino acid region (1–67) were synthesized and separately subcloned into the pUAST vector. The transgenic flies were then created via micro-injection at the Core Technology Facility of the Center for Excellence in Molecular Cell Science, Chinese Academy of Sciences (CAS). To simultaneously overexpress of *hTMEM41B* or *hTMEM41B*^ΔN^ while knocking down *stasimon* in muscles, *Stas^RNAi^* >*mef2-GAL4^ts^* flies were crossed with *hTMEM41H* or *hTMEM41B*^Δ*N*^ flies. Crosses were maintained at 29L. These strategies provided an effective system to study the functional interplay between TMEM41B variants and stasimon in muscle tissues.

### Asian black bear muscle tissue

Skeletal muscle tissues were obtained from forfeited frozen Asian black bears carcasses (*Ursus thibetanus*) provided by the Wildlife Forensic Institute of Northeast Forestry University. Sample collection and handling were conducted in accordance with national regulations governing the protection and rational utilization of wildlife resources. All procedures complied with relevant institutional and governmental guidelines and were approved by the appropriate regulatory authorities. Tissues were collected from the carcasses, snap-frozen in liquid nitrogen, and stored at –80 °C until analysis.

### Blue fox muscle tissue collection

Skeletal muscle samples were obtained from two healthy adult blue foxes (*Vulpes lagopus*) provided by a licensed fox breeding facility. Muscle tissues were immediately excised postmortem, snap-frozen in liquid nitrogen, and stored at -80L until further analysis.

### Cell culture and treatment

#### Myoblast isolation, C_2_C_12_ culture and myotube differentiation. Myoblast isolation

Tibialis anterior muscles were collected and digested with Collagenase D and dispase type II at 37L for 30 min. The digested tissue was filtered through a 70 μm cell strainer, and the filtrate was neutralized with 20-30 ml of DMEM. The cell suspension was centrifuged at 400 x g for 10 min at 4L, and the pellet was resuspended in growth medium (DMEM supplemented with 10% fetal bovine serum). After 2 hours, non-adherent cells were collected, resuspended in fresh growth medium, and cultured. Growth medium was replaced every two days (*41*).

#### C_2_C_12_ Myoblast Culture and Myotube Differentiation

C_2_C_12_ cells were maintained in growth medium as proliferating myoblasts. To induce myotube differentiation, C_2_C_12_ cells and primary myoblasts were cultured in an induction medium (DMEM supplemented with 2% horse serum). The induction medium was replaced every two days.

### TMEM41B stable knockout cell line construction

TMEM41B knockout cell lines were generated using CRISPR-Cas9 technology. Single guide RNA (sgRNA) sequences targeting Tmem41b were designed based on a previous publication (*42*) and cloned into the lentiCRISPR v2 vector (Addgene #52961). The resulting plasmids were transfected into C_2_C_12_ cells using lipo3000 following the manufacturer’s protocol. After 48 hours, transfected cells were subjected to puromycin selection until all non-transfected cells were eliminated. Surviving cells were selected with in growth medium (DMEM with 10% fetal bovine serum) and single-cell clones were established by serial dilution. Clonal cell lines were expanded and validated for downstream experiments. This approach enabled the efficient generation of stable TMEM41B knockout C_2_C_12_ cell lines for functional studies.

### HEK293T cell

HEK293T cells were obtained from ATCC and maintained in growth medium (High Glucose DMEM supplemented with 10% fetal bovine serum) under standard culture conditions. For transfection, cells were transfected with the indicated plasmids using polyethylenimine (PEI) following the manufacturer’s protocol.

### DNA vector construction

pCDNA3.1-CMV-51aa-TMEM41B-3xFLAG-hGHpolya-EF1a-EGFP (Bear-TMEM41B-GFP) and pCDNA3.1-CMV-TMEM41B-3xFLAG-hGHpolya-EF1a-EGFP(Human-TMEM41B-GFP) were purchased from OBiO Technology. The pLV2-CMV-V5-TurboID-Puro vector was purchased from MIAOLING BIOLOGY. Using this backbone, we constructed the following plasmids: pLV2-CMV-V5-TurboID-HumanTMEM41B (Human-TMEM41B), pLV2-CMV-V5-TurboID-BearTMEM41B(Bear-TMEM41B). These plasmids were transfected into HEK293T and C2C12 cells using polyethylenimine (PEI) or lipo3000, following the manufacturer’s protocol(*43*). Primers used were listed as follows: Xho1-human41B-F: GCCACTCGAGGCCACCATGGCGAAAGG; EcoR1-41B-R: GAACGAATTCCTCAAATTTCTGCTTTAGTTTTTTTTG; Xho1-51aa41B-F: GAACCTCGAGgccaccatggggagg.

### Body temperature under acute cold exposure

Cold-tolerance test was conducted both in 8- to 10-week and 7- to 10-day old mice. For ten-week-old mice, mice were housed individually at 4L with free access to food and water. Rectal temperature was measured using an electronic thermometer (Han Tai technology, TH212) before and after cold exposure. For 7- to 10-day old mice, body temperature was recorded before and after 0.5-to-1-hour cold exposure.

### Body temperature after anesthesia

Fourteen-week-old mice were anesthetized, and body temperature was measured using an electronic animal thermometer at baseline (pre-anesthesia) and at hourly intervals post-anesthesia. An Infrared digital thermography camera (InfraTec, VarioCAM HD head 980) was employed to monitor surface temperature changes. **Wake-Up Time.** The recovery time from anaesthesia to regained consciousness was also recorded.

### Surgical implanation of anipill capsules

Core body temperature was monitored in mice using an implantable biotelemetry system (Anilogger, BodyCAP). Anipill capsules were surgically implanted, activated, and paired with a monitor set to a specific radio channel. Data were recorded at defined intervals (e.g., 1-5 min), with simultaneous real-time transmission to the monitor and internal storage within the pill to ensure data integrity. The complete dataset was periodically offloaded via Anilogger Manager software for analysis.

### Body composition

Body composition in mice was conducted using a dual-energy X-ray absorptiometry (DXA) scanner, following the manufacturer’s guidelines. Fat mass and lean mass were quantified and analyzed to evaluate differences between experimental groups.

### Torpor induction and metabolic monitoring

*TMEM41B^flox/flox^* and *TMEM41B^Myl1-cre^* mice overespression either AAV-B41B or AAV-Con in muscles were singly housed in TSE Phenomaster animal monitoring system.After a accilimatization period at 25L with *ad libitum* access to food and water, mice were subjected to a 24-hour fast to induce torpor. Subsequently, mice were refed for a recovery peroid. Body temperature and energy metabolism was continuously throught the experiment.

### Micro-Computed Tomography (Micro-CT) analysis

Bone specimens (including femur and tibia) were dissected, fixed in 4% paraformaldehyde for 24 hours, and scanned using a high-resolution micro-CT system (SkyScan 1276, Bruker) at 60 kV/100 μA with a 10.0 μm isotropic voxel size. Reconstructed images were calibrated against a hydroxyapatite phantom for bone mineral density (BMD) and oriented along the long axis. Trabecular bone parameters were quantified in a 2.0 mm region of the distal femoral metaphysis, while cortical bone was assessed at the femoral mid-diaphysis over a 1.0 mm segment using CTan software. Three-dimensional visualizations were generated in CTvox.

### Chill coma recovery

Vials containing flies each were transferred from food vials to empty vials 10 min prior to cold exposure to acclimate. The vials were then submerged in ice water maintained at 0L for 20 min. Following cold exposure, the vials were immediately placed on a countertop at 25L for recovery. Recovery time was recorded using a timer, starting when all vials were placed on a white sheet of paper. Each vial was scored sequentially, and recovery for individual flies was defined as the time when a fly stood upright on all six legs (*44*).

### Bulk RNA Data Collection and Differential Analysis

First, we searched the GEO database for bulk RNA-seq datasets related to cold exposure in mice. A total of 41 research projects were curated and processed using in-house R (version 4.4.1) scripts, and differential expression gene (DEG) analysis was performed using the edgeR package (45) (version 4.2.2) after transferring all gene expression data back to raw counts. DEGs were identified based on Benjamini-Hochberg adjusted *p*-valuesL≤L0.05 using negative binomial distributions. We obtained 238 treatment-versus-control experiments in total and further counted the number of upregulated and downregulated genes in each experiment. Next, we compiled a list of all genes identified in the DEG files, sorting them in descending order based on the frequency of their differential expression for both upregulation and downregulation patterns across the datasets. We found that most thermogenesis-related genes (including Prkab2, Prdm16, Dio2, Ucp3, Ppargc1a, Cidea, Ucp1, Atp2a1, Actn3, and others) accounted for the top 5% of occurrences across the datasets. Therefore, the threshold for the so-called thermogenesis-related candidate genes was arbitrarily set at the top 5%.

### Gene Gain/Loss Analysis and Clustering

We aim to explore gene gain/loss analysis for thermogenesis-related candidates across species. Ensembl Homology REST API (https://rest.ensembl.org/documentation/info/homology_symbol) was used to fetch the homology relations of genes among 200 species. From the aforementioned DEG analysis, all candidate genes were matched with their corresponding cross-species homologous information, which we classified into 12 evolutionary clades of thermogenic mechanisms: Fungi, Nematoda, Arthropoda, Tunicata, Fish, Amphibia, Reptilia, Aves, Prototheria, Metatheria, small-size Mammals (s) and large-size Mammals (l). Next, we constructed a ‘gene loss vector’ to assess the similarity of gene gain/loss patterns between any two genes. For each evolutionary clade, we classified a gene as either present (with one or multiple copies) or absent (zero copies) in each species and gathered the genes within the clade. Thus, for a strict single-copy gene across all clades, the two-dimensional nested vector would, in principle, take the form ((1…1), (1…1), …, (1…1)), representing the presence of the gene across all species in clades. To represent the evolutionary relationships among these clades, we constructed a phylogenetic tree using iTOL (v7) (*46*). Gene gain was not weighted above 1, as thermogenesis-related genes typically lack paralogs (except in fish, which experienced additional whole-genome duplication events). Therefore, we hypothesize that the expansion of genes into multiple copies is not a common mechanism underlying the evolution of thermogenesis and was excluded from consideration in our study.

Additionally, we merged the species from Fungi, Nematoda, Arthropoda, and invertebrate Chordates as an outgroup of vertebrates and excluded clades with fewer than ten species (Amphibia, n = 2; Monotremata, n = 1; Marsupialia, n = 5) during the clustering process. These clades were recalled only for manual assessment after clustering. Notably, although some fish species have developed regional endothermy (*^47^*), they are not recorded in the current Ensembl database. Therefore, the fish clade could not be further partitioned based on thermogenic mechanisms. The final vector *v* was constructed as the gene copy numbers across the following clades: (Fish, Reptilia, Aves, Mammal (s), Mammal (l)).

The distance metric between each pair of genes was assessed using:

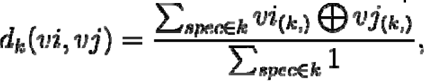

for gene *i* and *j* across every species *spec* in clade *xpec*, where (Fish, Reptilia, Aves, Mammal (s), Mammal (l)). LJ denotes ’Exclusive OR’, with the following operational rules: 0 LJ 0 = 0, 1 LJ 0 = 1, 0 LJ 1 = 1, and 1 LJ 1 = 0. The metric is analogous to the Jaccard distance for assessing beta diversity, while accounting for both the co-occurrence and co-absence of gene pairs within the same species.

To verify the results and interpret/annotate the clusters, we combined all potential clade-wise gene loss events based on the thermogenesis tree to generate a set of pseudogenes as spike-ins. For example, a pseudogene that loses all gene copies in Reptilia and Aves was represented as (1_*n*1_, 0_*n*2_, 0_*n*3_, 1_*1*n1_, 1_*n*5_) (*n* is the total number of the species in the corresponding clade) and, in principle, should be located very close to UCP1.

Pseudogenes that were far from the bona fide genes were defined as evolutionary events that did not actually occur and were therefore excluded from the final representation. The clusters were ultimately annotated based on the pseudogene closest to the cluster centroid(*48*).

### Generation of a Candidate NST Gene Set

We curated 9 canonical NST genes and 9 pseudogenes (after excluding the ‘loss in all’ pseudogene) based on the UMAP visualization as anchors. Next, we selected the 30 nearest-neighbor genes (including ties) for each anchor, thereby subsetting the thermogenesis-related candidate DEGs into a smaller set for downstream sequence conservation analysis. The gene list is provided in Supplementary Table S2.

### Measurement of Sequence Conservation

First, the phastCons score (*22*) file for the human genome (GRCh38) was downloaded from UCSC Genome Browser (https://hgdownload.soe.ucsc.edu/goldenPath/hg38/phastCons470way/hg38.phastCons470way.bw). The Ensembl canonical transcript was used to represent the typical transcript of each gene, and their genome coordinations were extracted from the Ensembl annotation file. (https://ftp.ensembl.org/pub/current_gtf/homo_sapiens/Homo_sapiens.GRCh38.113.gtf.gz). It is noticed that in a large proportion of our interest genes, the phastCons score varies significantly on different regions of the gene, typically with the 3’-end or 5’-end less conservative and the middle of the gene more conservative. To address this issue, the CDS regions were split into 45-nucleotide (equivalent to 15 amino acids) blocks for each gene, and the average phastCons score was calculated over each block. The median of block-averaged scores was further calculated for each gene and ranked among all genes to represent its conservation status. In the box plot representation, the median score (indicating overall sequence conservation), the lower whisker, and the lower outliers (highlighting locally evolved domains) were of particular interest.

### N-terminal plasticity in sequences and protein structures

The N-terminal region of *TMEM41B* was defined as 1∼67 amino acids(aa) in the human protein, where the 67 aa is preceding the nearest start codon of the first transmembrane structure. In other species, the N-terminal boundaries were determined based on pairwise alignments to the human protein.

To obtain homologous sequences for comparative genomics, genome assemblies from 82 representative Boreoeutherian mammals spanning both Euarchontoglires and Laurasiatheria (table S3). Each genome was formatted as a searchable BLAST database using the makeblastdb(v2.17.0+) (*-dbtype nucl - parse_seqids*)(*49*).

*TMEM41B* homologs were identified using the BLASTN algorithm (*-task dc-megablast -evalue 1e−3 -word_size 7 -reward 2 -penalty −3 -gapopen 5 -gapextend 2*), and the top hit with the highest bitscore for each query was retained as the best putative ortholog for downstream analysis. Gene structures of *TMEM41B* orthologs were visualized using the R packages gggenes (*50*) and ggplot2 (*51*), based on annotated exon–intron boundaries and UTRs.

To quantify evolutionary divergence, normalized Hamming distances were calculated from the pairwise alignment results for N-terminal and rest region respectively, and normalized by dividing the corresponding pairwise alignment length.For broader N-terminal comparisons, *TMEM41B* homologs from representative mammalian families were aligned using MUSCLE5 (v5.3)(*52*) for multiple sequence alignment (MSA). For MSA loci covered by more than 10% sequences, the sequence consistency was calculated by adding the pairwise similarity score from BLOSUM62 matrix, and normalizing into range [-1,1] by dividing the maximum possible score in theory. The consistency score is calculated using a modified version of trimAl (*^53^*). Sequence alignments at both nucleotide and amino acid levels were visualized using the R package ggmsa (*54*).

The protein structures are predicted by AlphaFold (v1.01)(*55*) in UCSF ChimeraX (v1.9) (*56*)(2024-12-11). All animals silhouette images used in the figures were sourced from PhyloPic (http://phylopic.org) under a public domain license.

### Phylogenetic Analysis

All phylogenetic analyses of sequence data were conducted using IQ-TREE (v3.0.1)(*57*), with the best-fit substitution model selected by ModelFinder under the Bayesian Information Criterion (BIC) *(-m MFP)*. Node support assessed using 1,000 ultrafast bootstrap replicates incorporating the NNI optimization step *(-B 1000 -bnni)*. In addition, the species phylogeny was constructed and modified based on the TimeTree5(*58*) to reflect the taxonomic relationships among the analyzed taxa.

### Transcriptomic Processing and N-terminal Coverage Analysis of *Tmem41b* in Bears

We retrieved publicly available bear transcriptomic study and datasets from the NCBI SRA database, including PRJNA747307, PRJNA605162, PRJNA514749, and PRJNA1011923, corresponding to the polar bear (*Ursus maritimus*), American black bear (*U. americanus*), brown bear (*U. arctos*), and giant panda (*Ailuropoda melanoleuca*), respectively (*59–61*) (table S4). The polar bear dataset comprised ONT-based R2C2 full-length cDNA sequencing, while the remaining datasets represented illumina-based short-read sequencing data.

Polar bear raw reads were processed with C3POa (v3.1)(*62*) (https://github.com/rvolden/C3POa) to generate high-accuracy R2C2 consensus sequences. The resulting consensus reads were mapped to the U. maritimus reference genome (GCF_017311325.1) using minimap2 (v2.26) (*63*) with spliced alignment parameters (*-ax splice -uf -k14*). Illumina reads were quality controlled and adapter-trimmed with Trimmomatic (v0.40) (*64*) using the following parameters: *ILLUMINACLIP:$ADAPTERS:2:30:10 LEADING:3 TRAILING:3 SLIDINGWINDOW:5:20 MINLEN:36 HEADCROP:10*. Clean reads were then aligned to the corresponding reference genomes from NCBI RefSeq (giant panda: GCF_002007445.2; American black bear: GCF_020975775.1; brown bear: GCF_023065955.2) by building genome indices and mapping using HISAT2 (v2.2.1) (*65*) with default parameters (*hisat2 -build*; *hisat2 -S*).

We used SAMtools (v1.16.1) (*65*) to converted all alignment files from SAM to BAM format, sorted, and indexed (*samtools sort*; *samtools index*). The N-terminal region of *TMEM41B* was extracted using *samtools view -b*, and coverage depth across this region was visualized using the IGV software (v2.19.5) (*66*).

### Hematoxylin and Eosin (H&E) Staining

Muscles were mounted on cork using gum tragacanth and frozen in liquid nitrogen-cooled isopentane. Brown adipose tissue (BAT), inguinal white adipose tissue (iWAT) and epididymal white adipose tissue (eWAT) were fixed in 4% paraformaldehyde for 24 hours and subsequently embedded in paraffin. Paraffin-embedded tissues were sectioned at a thickness of 5 μm, while cryo-embedded tissues were sectioned at 10 μm. Sections were then subjected to hematoxylin (E607317-0500, Sangon Biotech) and eosin (E607321-0100, Sangon Biotech) following standard protocols.

### Transmission electron microscopy and analysis

Tibialis anterior muscles and BAT tissues were collected and pre-fixed in glutaraldehyde (Solarbio, P1126) at 4L. Tissues were then processed, sectioned, and stained according to standard protocols. Ultra-thin sections were imaged using a JEM-2100 transmission electron microscope (Hitachi HT7700, Japan) (*67*). The spatial relationship of mitochondria and endoplasmic reticulum (ER) were generated from a DeepContact model (*68*).

### Co-localization analysis

HEK293T cell and C_2_C_12_ cells were cultured in Dulbecco’s modified Eagle’s medium (DMEM) with 10% fetal bovine serum (FBS) at 37°C with 5% CO_2_. Human TMEM41B-Flag plasmid and Bear-TMEM41B-Flag plasmid were co-transfected with ER-DsRed and Mito-GFP respectively. Cells were transfected with indicated plasmids using polyethylenimine (Lipo3000, thermo fisher) according to the instructions. After 48 hours, cells were harvested for immunofluorescence staining. The primary antibody was Flag (Abclonal, AE092), with secondary antibodies comprising Alexa Fluor 488 anti-rabbit IgG (H+L) (Thermo Fisher Scientific, A-11001) and Alexa Fluor 594 anti-rabbit IgG (H+L) (Thermo Fisher Scientific, A-11005). Imaging was performed using a Leica confocal microscope, and quantitative analyses were carried out using Fiji software.

### Mitotrakcer and JC-1 imaging

To assess mitochondrial membrane potential and superoxide anion levels, primary muscle cells or C_2_C_12_ cells were seeded onto confocal culture dishes. After cell adhesion, staining was performed using the JC-1 Mitochondrial Membrane Potential Assay Kit (Solarbio, M8650), and MitoTracker (Invitrogen, M22425). Imaging was conducted using a Leica confocal microscope.

### Mitochondrial respiration studies

Muscle permeabilization was performed as previously described (*67*). For brown adipose tissue mitochondria exposure: approximately 2 mg tissue was homogenized in MIR05 buffer (0.5 mM EGTA, 3 mM MgCl_2_, 60 mM lactobionic acid, 20 mM taurine, 10 mM KH_2_PO_4_, 20 mM HEPES, and 110 mM sucrose). For myotube permeabilization, a total of 1 x 10^6^ cells were transferred into the O_2_K chambers and permeabilized with 5 ul digitonin (10 mg/ml). 5 mM pyruvate, 2 mM malate, 10 mM glutamate, 2.5 mM ADP and 10 µM cytochrome C were used to measure the complex I respiration flux; 1 M succinate was added for complex II; 0.5 µM step carbonyl cyanide 4-(trifluoromethoxy) phenylhydrazone was added until maximal respiration flux; 0.5 µM rotenone was added for respiration through complex I; 2.5 µM antimycin was added for ending the measurement.

### Protein extraction and Immunoblotting

Muscles, BAT, iWAT, heart and cultured cells were lysed in RIPA buffer (Beyotime, P0013B) supplemented with PMSF (Beyotime, P1006). Protein concentrations were determined using a BCA assay kit (Beyotime, P0010). Equal amounts of protein were resolved on SDS-PAGE gels and transferred onto PVDF membranes (Millipore). Then membranes were blocked with 5% non-fat milk and incubated overnight at 4L with primary antibodies. After washing with 1×TBST, membranes were incubated with HRP-conjugated secondary antibodies. Protein signals were detected using an ECL substrate (LINDE sentific, LP2003), and visualized using a chemiluminescence imaging system (SAGECREATION). Densitometry analysis was conducted using ImageJ software. The primary antibodies used in the current study were: SDHB (Proteintech, Cat no. 10620-1-AP, RRID: AB_2285522), NDUFB8 (Proteintech, Cat no. 14794-1-AP, RRID: AB_2150970), UCP1 (Abcam, Cat no. ab10983, RRID:AB_2241462), VDAC1 (Abclonal, Cat no. A19707, RRID: AB_2862746), LC3 (MBL, Cat no. PM036, RRID: AB_2274121), P62 (Proteintech, Cat no. 18420-1-AP, RRID: AB_10694431), PARKIN (Abclonal, Cat no. A0968, RRID: AB_2757487), TMEM41B (Proteintech, Cat no. 29270-1-AP, RRID: AB_2918264), V5 (Cell signaling technology, Cat no. 13202, RRID: AB_2687461), Flag (Abclonal, Cat no. AE005, RRID: AB_2770401), GFP (Abclonal, Cat no. AE078, RRID: AB_3711256), Myc (Abclonal, Cat no. AE070, RRID: AB_2863795), GAPDH (Proteintech, Cat no. 60004-Ig, RRID: AB_2107436), β-actin (Abclonal, Cat no. AC026, RRID: AB_2768234), Tubulin (Abclonal, Cat no. A12289, RRID: AB_2861647).

### CO-Immunoprecipitation (CO-IP)

For immunoprecipitation(*69*), cells were lysed in RIPA buffer (50Lmmol/L Tris-HCl (pH 7.4), 150Lmmol/L NaCl, 1% sodium deoxycholate, 1% Triton-100, plus 0.1% SDS, Beyotime, P0013B) containing complete Protease Inhibitor Cocktail (Beyotime, P1005) and centrifuged at 12,000 g for 10 min at 4 °C. A portion of the supernatant was mixed with 5x loading buffer to serve as the input sample. The remaining lysates were incubated with Flag Magarose beads (AlpaLifeBio, KTSM2425, avoid IgG) at 4 °C overnight. After washing the beads three times in RIPA, they were boiled with 5x loading buffer for 10 min and then subjected to western blot analysis using the indicated antibodies.

### RNA extraction and RT-qPCR

Total RNA was extracted from tissues or cells using TRIzol Reagent (Invitrogen) following the manufacturer’s instructions. RNA concentration was detected using a NanoDrop spectrophotometer (Thermo Fisher Scientific). Complementary DNA (cDNA) was synthesized using the High-Capacity cDNA reverse transcription kit (Applied Biosystems). Quantitative PCR (qPCR) was performed using SYBR Green Master Mix (Applied Biosystems) on a QuantStudio 5 Real-Time PCR System (Applied Biosystems). Relative gene expression levels were calculated using the 2^−ΔΔCt^ method, with *Tbp* or *36b4* serving as the internal reference. Primers used in qPCR were listed as followed: *Cidea*, TGCTCTTCTGTATCGCCCAGT and GCCGTGTTAAGGAATCTGCTG; *Dio*2, TCCTAGATGCCTACAAACAGGTTA and GGTCAGGTGGCTGAACCAAA; *Ucp1*, ACTGCCACACCTCCAGTCATT and CTTTGCCTCACTCAGGATTGG; *Rel*, CTCTGCCTCCCATTGTTTCTA and GGCTTCCCAGTCATTCAACAC; *RelA*, GCCCAGACCGCAGTATCC and GTCCCGCACTGTCACCTG; *RelB*, CTGGCTCCCTGAAGAACC and CGCTCTCCTTGTTGATTC; *Il-10*, GCTCTTACTGACTGGCATGAG and CGCAGCTCTAGGAGCATGTG; *Il-1*β, GAAATGCCACCTTTTGACAGTG and TGGATGCTCTCATCAGGACAG; *Tnf*α, CAGCGCTGAGGTCAATCTGCC and TGCCCGGACTCCGCAA; *Tgf*β, TTGCTTCAGCTCCACAGAGA and TGGTTGTAGAGGGCAAGGAC; *Inos*, GTTCTCAGCCCAACAATACAAGA and GTGGACGGGTCGATGTCAC; *Arg1*, AAAAGCCAAACTATTGCCCAGA and TCCCACCATTCCTTTCCTTGC; *Il-6*, ATCCAGTTGCCTTCTTGGGACTGA and TAAGCCTCCGACTTGTGAAGTGGT; *Tmem41b*, AACATTTGCTATCCCAGGTTCC and ACGGGTCTTCCAACTAAGTAGG; *Tbp*, ACCCTTCACCAATGACTCCTATG and TGACTGCAGCAAATCGCTTGG; *36b4*, TGCTGAACATCTCCCCCTTCTC and TCTCCACAGACAATGCCAGGAC; *Stasimon*, GTCGCCTACGTATTCCTGCAAACA and CCAGCGCCGAACAGAAACATATGA; *Rp49*, GCAAGCCCAAGGGTATCGA and TAACCGATGTTGGGCATCAG; Bear-*Tmem41b*,GGCGGAGTCTAGTCCGAGTG and TCGTTCGGCGACTCTGC.

### PCR of TMEM41B in bear and blue fox muscle

After reverse transcripiton of bear and fox muscles’ RNA as described, we detected the N terminal of TMEM41B by PCR and then ran 1.5% agrose gel in 1X TAE buffer 140V 20min. Taq mix was purchased from Abclonal(RK20718) and the PCR programme was used from its product manual. Primers used were listed as follows: Bear41B-N-F: GACCCTGACACGTCTCGC; Bear41B-N-R: TGACATTCTTGCTGACCCA; Bear41B-C-F: AGGTTGAGCGTCATAGA; Bear41B-C-R: AAGATGGCTGGCAGAATAGAAAGAA; Fox41B-N-F: gaccctgacttgtctcgc; Fox41B-N-R: TGACATTCTTGCTGACCCA; Fox41B-C-F: AGGTTGAGCGTCATAGA; Fox41B-C-R: aagatggctggcaggatagaaagaa.

### DNA Extraction and PCR Amplification of *Drosophila*

*Drosophila* were anesthetized and lysed in 1× Mouse Tissue Lysis Buffer (Vazyme) containing Proteinase K. Samples were incubated at 55 °C for 45 min, followed by 95 °C for 5 min. Lysates were centrifuged at 15,000 rpm for 5 min at 4°C. DNA was extracted using an equal volume of extraction buffer, precipitated with isopropanol, washed with 75% ethanol, and dissolved in distilled water. DNA concentrations were determined spectrophotometrically. Specific primers for full-length and truncated target genes were designed and synthesized (Sangon Biotech, Shanghai, China). Primer sequences included: primer1, GAACGAATTCATGGCGAAAGGCAGAGTC and GAACCTCGAGCTCAAATTTCTGCTTTAGTTTTTTTTGGAAG; primer 2, GAACGAATTCATGTTTTTGGTATATAAAAATTTTCCTCAGCTT and GAACCTCGAGCTCAAATTTCTGCTTTAGTTTTTTTTGGAAG.PCR products were analyzed on 1% agarose gels. Bands were visualized under UV light using an automatic gel imaging system and compared to a 100 bp DNA Ladder to confirm the expected fragment sizes.

### RNA-sequencing

BAT and muscle tissues from each group were collected for RNA sequencing. Total RNA was extracted using the Trizol method according to the manufacturer’s instructions. RNA quality was evaluated with an Agilent 2100 Bioanalyzer using the RNA 6000 Nano Kit from Agilent Technologies (Santa Clara, CA, USA). cDNA libraries were prepared for each sample following the previously established protocols (*70*). Raw sequencing reads were filtered using Trim Galore (v0.6.7) to remove adapters and low-quality bases. The resulting clean reads were aligned to the mouse reference genome (mm10) using STAR (v2.5.2b). Gene expression levels were quantified with RSEM (v1.2.28) using the following command “rsem-calculate-expression --paired-end -p 20 --alignments samples. bam star_index.files”. Differential gene expression between *Tmem41b* knockout (*Tmem41b* AKO) and wide type (*TMEM41B^f/f^*) groups, *Tmem41b* knockout (*Tmem41b* MKO) and wide type (*TMEM41B^f/f^*) groups was analyzed using DESeq_2_ (v1.42.0), Genes with an adjusted *p* value < 0.05 and |fold change| > 1.5 as significantly differentially expressed. The significantly differentially expressed genes were subjected to KEGG pathway enrichment analysis to identify functional pathways associated with *Tmem41b* knockout.

### Overexpression of human TMEM41B or bear TMEM41B in C2C12 cells, cell sorting and RNA sequencing

C_2_C_12_ myoblasts were transfected with plasmids expressing human-TMEM41B-GFP or a Bear-TMEM41B-GFP using lipo3000 according to the manufacturer’s instructions. Forty-eight hours after transfection, GFP-positive cells were isolated by fluorescence-activated cell sorting (FACS) to ensure enrichment of successfully transfected populations. Sorted cells were collected, and total RNA was extracted using TRIzol reagent (Invitrogen) following the manufacturer’s protocol. RNA quality and integrity were assessed prior to library preparation, and high-quality samples were subjected to RNA sequencing for transcriptomic analysis.

### Trypsin digestion

The FASP digestion protocol was adapted using Microcon PL-10 filters, following previously described methods (*71*). Proteins were subjected to three buffer exchanges with 8 M urea and 100 mM Tris-HCl (pH 8.0). Reduction was performed with 10 mM dithiothreitol (DTT) at 37L for 30 min, followed by alkylation with 30 mM iodoacetamide at 25L for 45 min in the dark. Subsequently, the buffer was exchanged three times with a digestion buffer (30 mM Tris-HCl, pH 8.0). Proteolytic digestion was conducted with trypsin at an enzyme-to-protein ratio of 1:50 at 37L for 12 hours. After digestion, the resulting solution was filtered, and the filter was washed twice with 15% acetonitrile (ACN). All filtrates were pooled, vacuum-dried, and stored for subsequent analysis.

### Overexpression of human TMEM41B or bear TMEM41B and immunoprecipitation-mass spectrometry

The pLV2-CMV-HumanTMEM41B-V5-TurboID (Human-TMEM41B), pLV2-CMV-Bear51aaHumanTMEM41B-V5-TurboID (Bear-TMEM41B) or pLV2-CMV-V5-TurboID (con) were transfected into HEK293T using polyethylenimine (PEI). Forty-seven hours after transfection, 100uM biotin was added to cell dishes separately. Forty-eight hours after transfection, cells were lysed in RIPA buffer (Beyotime, P0013B) supplemented with PMSF (Beyotime, P1006). The lysates were cleared by centrifugation, and then the supernatants were treated following a previously described protocol(*43*). Bound proteins were eluted, resolved by SDS-PAGE and sliver staining (Beyotime, P0017S), then subjected to in-gel digestion. LC-MS analysis was conducted using an EASYnLC 1200 nanoflow system (Thermo Fisher Scientific, Odense, Denmark) interfaced with an Orbitrap Exploris 480 mass spectrometer (Thermo Fisher Scientific, Bremen, Germany). A single-column configuration was employed for all analyses. Peptides were separated on a custom-packed C18 analytical column (75 µm inner diameter × 25 cm, ReproSil-Pur 120 C18-AQ, 1.9 µm; Dr. Maisch GmbH, Germany) (*72*). The mobile phases consisted of Solution A (0.1% formic acid) and Solution B (0.1% formic acid in 80% acetonitrile). For LFQ analysis, peptides were eluted using the following gradient: 5–8% B over 3 min, 8–44% B over 100 min, 44–70% B over 5 min, 70–100% B over 2 min, followed by 10 min at 100% B, at a flow rate of 200 nL/min. High-field asymmetric waveform ion mobility spectrometry (FAIMS) was activated during data acquisition with compensation voltages set at −40 and −60 V. MS1 spectra were acquired in the Orbitrap at a resolution of 60,000. Precursors with charge states between 2 and 7 were selected for MS2 analysis, with a dynamic exclusion window of 45 s. The cycle time was set to 1.5 s. MS2 spectra were collected in the Orbitrap using HCD fragmentation (isolation window 1.6; resolution 15,000; normalized collision energy 30%; maximum injection time 30 ms).

### Raw Data Processing

The resulting data were analyzed using the UniProt human protein database (75,004 entries, downloaded on 2020-07-01) and processed with Proteome Discoverer (version 2.4, Thermo Fisher Scientific) employing Mascot (version 2.7.0, Matrix Science) (*73*). Mass tolerances were set to 10 ppm for precursor ions and 0.05 Da for fragment ions. Up to two missed cleavages were permitted. Carbamidomethylation of cysteine was specified as a fixed modification, while acetylation of the protein N-terminus and oxidation of methionine were considered variable modifications. Gene Ontology (GO) analysis and the Kyoto Encyclopedia of Genes and Genomes (KEGG) analysis were performed using clusterProfiler v4.8.3 package in R (version 4.3.0). All the visualization methods of proteomics are based on dplyr and ggplot2 packages.

### Statistical analysis

Mice and flies were randomly assigned to experimental groups, and investigators were blinded to group allocation during data collection and analysis. Statistical analysis was performed using GraphPad Prism 9. Data are presented as mean ± s.e.m., with error bars representing the standard error of the mean. Group comparisons were performed using two-tailed unpaired Student’s *t*-test or one-way ANOVA followed by Bonferroni’s post-hoc test. For chill coma recovery experiments, the Log-rank (Mantel-Cox) test was applied. Statistical significance was defined as *p* < 0.05.

## Data availability

The RNA-seq data have been deposited in the GEO database under the accession numbers GSE287786 (TA) and GSE287714 (BAT), EMBL-EBI database under the accession numbers E-MTAB-16283 (C_2_C_12_ cell line). All of the other data have been deposited in the STOmics Database of the China National GeneBank Database (CNGBdb) under the accession number CNP0006837. A temporary link (https://db.cngb.org/search/project/CNP0006837/) is provided for open-access review at this stage.

## Code availability

Codes used in this study have been deposited at GitHub and can be accessed at https://github.com/BrainStOrmics/Thermogensis_Analyses.

## Acknowledgements

The authors acknowledge the assistance of the platform of State Key Laboratory of Genetics and Development of Complex Phenotypes at Fudan University. This work was funded by the Noncommunicable Chronic Diseases-National Science and Technology Major Project (2024ZD0530400), National Natural Science Foundation of China (82530026, 92457301, 92157203), Cao Ejiang Basic Research Fund of Fudan University (24FCB07), Shanghai Municipal Science and Technology Major Project,International Human Phenome Project II (2023SHZDZX02D24) to X.K., the National Key R&D Program of China (2025YFA1804600), the National Natural Science Foundation of China (82570994, 92357304), Faculty Resources Project of College of Life Sciences, Inner Mongolia University (2022–102), State key Laboratory of Reproductive Regulation & Breeding of Grassland Livestock (2025KYPT0081), Funding for the Flexible Talent Program of the Human Resources and Social Security Department of Jilin Province 2025 to T.L., the China Postdoctoral Science Foundation (2024M760516), the Postdoctoral Fellowship Program of CPSF under Grant Number (GZC20230527) to S.Z.,National Natural Science Foundation of China (32501027),the Postdoctoral Fellowship Program of CPSF under Grant Number (GZC20251821), the China Postdoctoral Science Foundation (2025M772758), the Noncommunicable Chronic Diseases-National Science and Technology Major Project (2025ZD0550000) to H.W., the Fund of Shanghai Municipal Health Commission (20204Y0246),Yangfan Project of Shanghai Science and Technology Commission (21YF1405000) and the National Natural Science Foundation of China (82300978) to Y.F..

This work is supported by the High-performance Computing Platform of Yazhou Bay Science and Technology City Advanced Computing Center (YZBSTCACC).

## Author contributions

X.K., Y.B. and S.F conceptualized the study. Methodology was developed by Y.X., J.Z., Y.F., H.W. and Y.W.. S.Z., X.Z. and P.Z.. conducted the investigation. Visualization was carried out by K.X., Y.L., Y.C., J.Y. and H.Z.. Funding acquisition was secured by X.K., T.L., S.Z., H.W. and Y.F.. Project administration was managed by T.L. X.L., and H.Y.. C.H and J.Y supervised the study. Y.M. and Y.X provide the samples. X.K. and Y.B. prepared the original draft, with subsequent review and editing by S.Z., H.W. and Y.Z..

## Supplementary figures

**Fig. S1.**
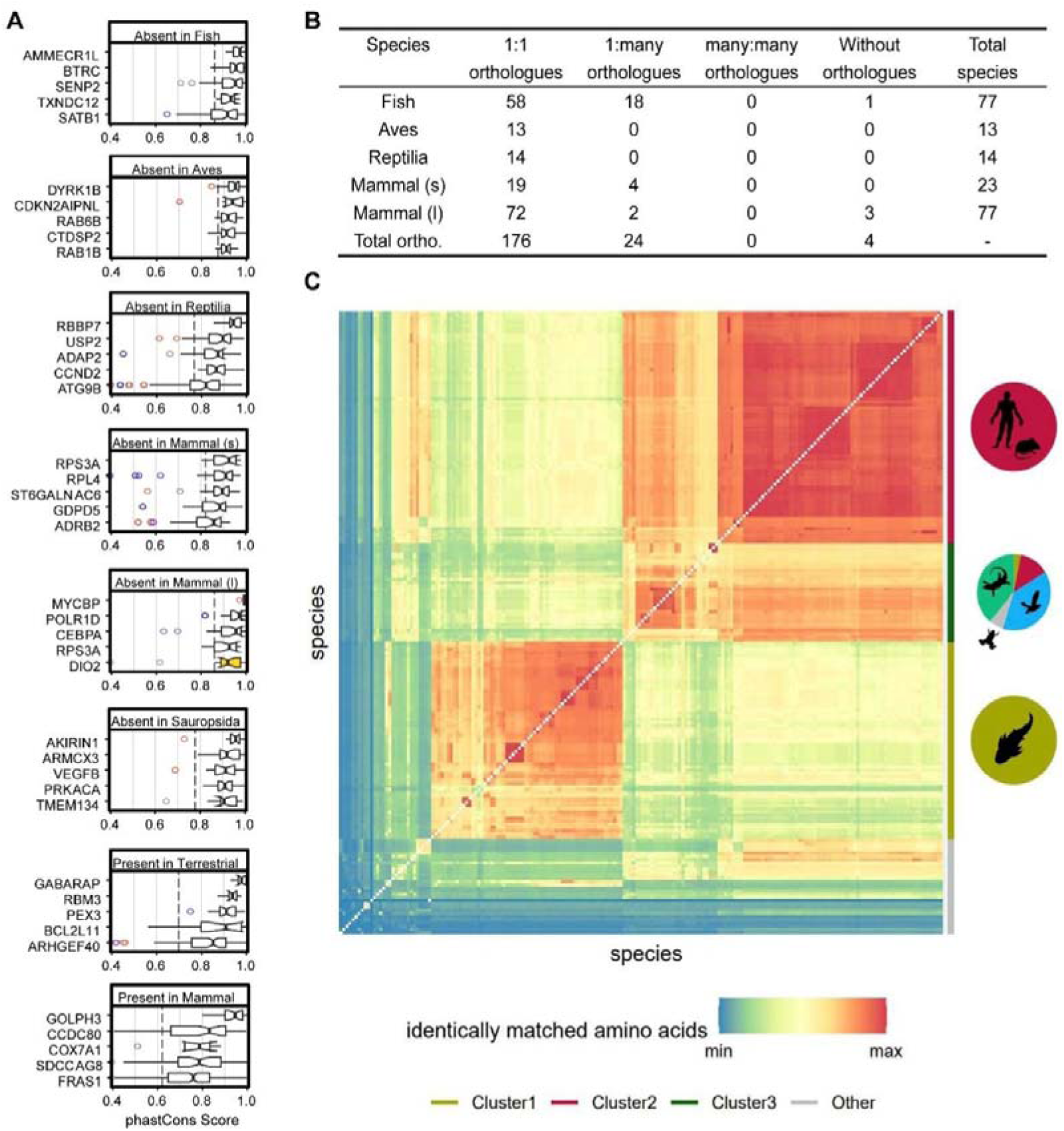
TMEM41B is an evolutionarily conserved single-copy gene while exhibiting rapid evolution from aquatic to terrestrial vertebrates. (**A**) Boxplots showing the top 5 conserved genes across other clusters, related to Figure 1E. (**B**) Summary of orthologues of TMEM41B. (**C**) Heatmap showing the hierarchical clustering of TMEM41B sequence homologies across all 196 species from the Ensembl database. *Aquatic* (bottom left) and *terrestrial* (upper right) vertebrates are distinctly separated.

**Fig. S2.**
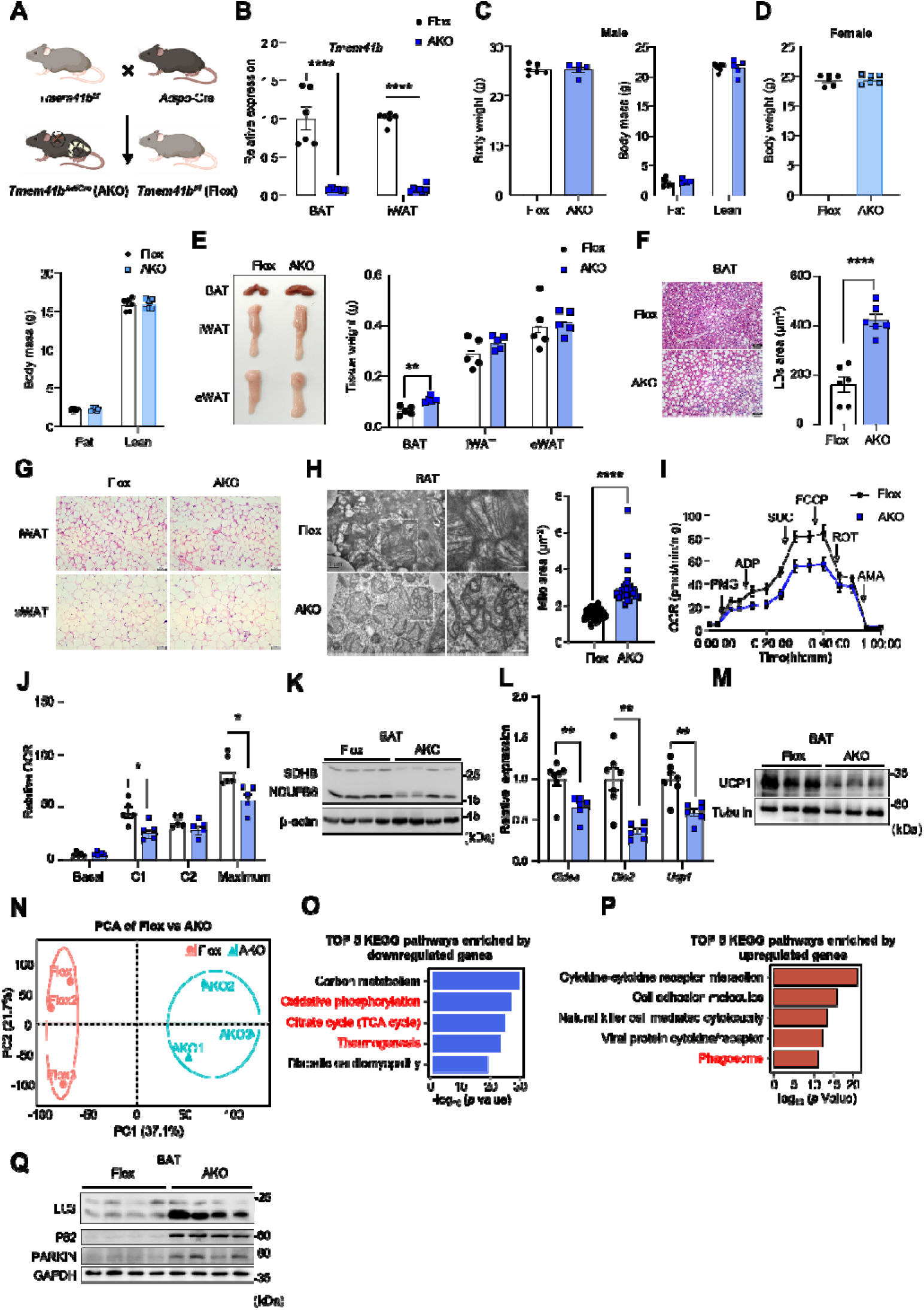
Impaired thermogenic response and mitochondrial function in adipocytes specific *Tmem41b* knockout mice. (**A**) Schematic diagram showing the strategy for generating adipose tissue-specific *Tmem41b* knockout (*Tmem41b^AdiCre^*) mice. (**B**) Relative mRNA expression levels of *Tmem41b* in BAT and iWAT tissues from *Tmem41b^f/f^* and *Tmem41b^AdiCre^* mice, normalized to *36b4* (n =6, 5 per group). (**C-D**) Body weight and body composition male and female mice (n = 6, 5 per group). (**E**) Representative images and tissue weights of brown adipose tissue (BAT), inguinal white adipose tissue (iWAT), and epididymal white adipose tissue (eWAT) from *Tmem41b^f/f^*and *Tmem41b^AdiCre^* mice (n = 5 per group). (**F-G**) Representative hematoxylin and eosin (H&E) staining of BAT,iWAT and eWAT from *Tmem41b^f/f^* and *Tmem41b^AdiCre^* mice. Scale bar = 50 µm. (F) BAT; (G) iWAT and eWAT. (**H**) Transmission electron microscopy (TEM) of BAT reveals enlarged mitochondria with disrupted cristae in *Tmem41b^AdiCre^* mice. Scale bar = 2 µm. Right, Mitochondrial area quantification. (**I**) Representative traces of oxygen consumption rates (OCR) in BAT from *Tmem41b^f/f^* and *Tmem41b^AdiCre^*mice (n = 5 per group). (**J**) Quantification of basal, complex 1 (C1), complex 2 (C2) and maximal respiratory capacity (Maximum) in BAT from *Tmem41b^f/f^*and *Tmem41b^AdiCre^* mice (n = 5 per group). (**K**) Western blot analysis of SDHB and NDUFB8 proteins in BAT (n = 4 per group). (**L**) RT-qPCR analysis of thermogenic genes in BAT (n = 7, 6 per group). (**M**) Expression levels of thermogenesis-related proteins (UCP1) in BAT (n = 3 per group). (**N**) Principal component analysis (PCA) of RNA-seq data of BAT from *Tmem41b^f/f^* and *Tmem41b^AdiCre^* mice (n = 3 per group). (**O**) Top five enriched KEGG pathways of downregulated genes in BAT of *Tmem41b^AdiCre^*mice in bulk RNA-seq analysis. (**P**) Top five enriched KEGG pathways of upregulated genes in BAT of *Tmem41b^AdiCre^* mice. (**Q**) Western blot analysis of autophagy-related proteins, including LC3, P62, and PARKIN, in BAT from *Tmem41b^AdiCre^* mice compared to *Tmem41b^f/f^* controls (n = 4 per group).

**Fig. S3.**
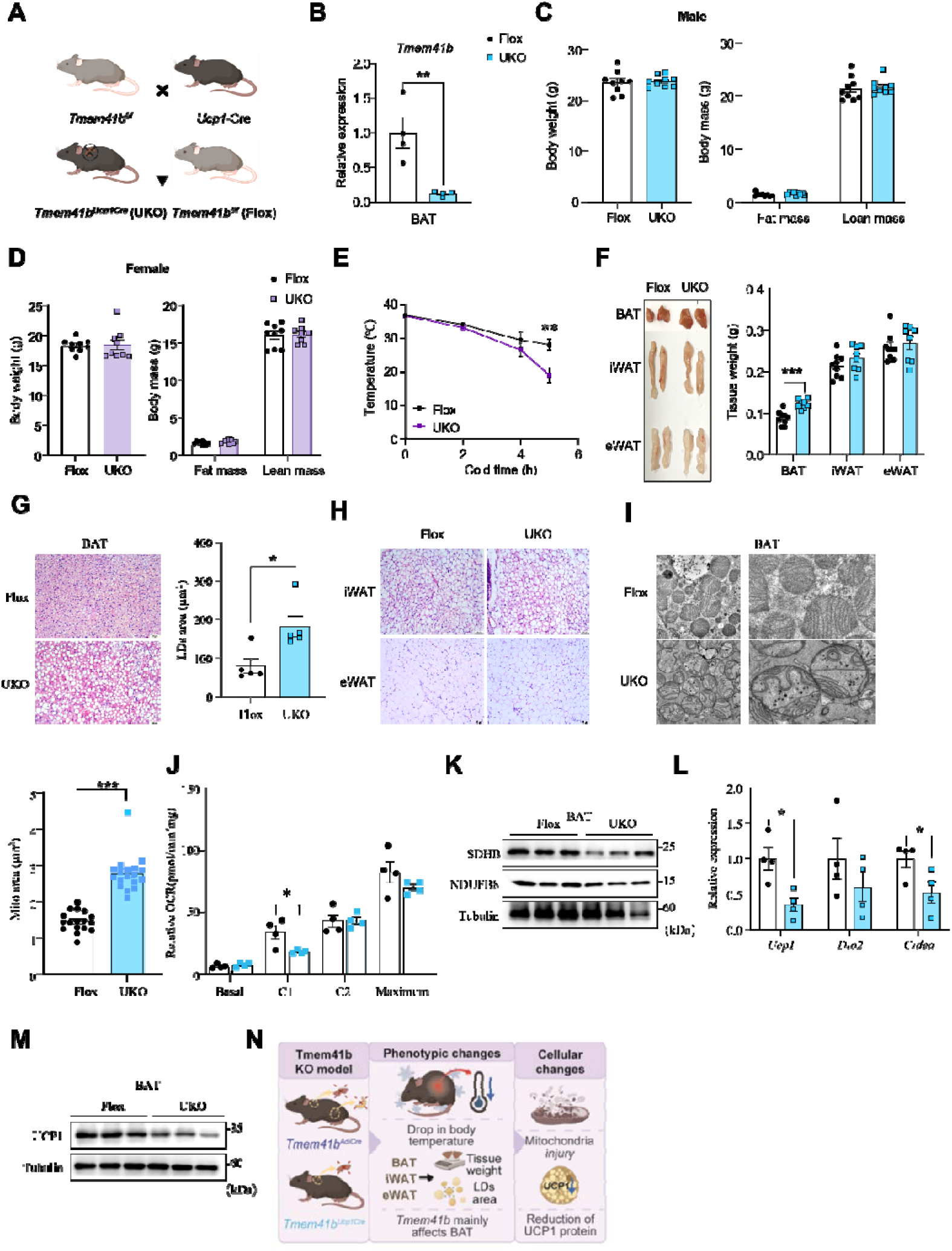
Impaired thermogenic response and mitochondrial function in brown adipocytes-specific Tmem41b knockout mice. (**A**) Schematic diagram showing the strategy for generating brown adipocyte-specific *Tmem41b* knockout (*Tmem41b^Ucp1Cre^*) mice. (**B**) Relative mRNA expression levels of *Tmem41b* in BAT tissue from *Tmem41b^f/f^* and *Tmem41b^Ucp1Cre^* mice, normalized to *36b4* (n = 4 per group). (**C-D**) Body weight and body composition of male and female mice (n = 9 per group). (**E**) Cold tolerance test of female mice (n = 6 per group). (**F**) Tissue weights and representative images of BAT, iWAT and eWAT from *Tmem41b^f/f^* and *Tmem41b^Ucp1Cre^* mice (n = 9, 8 per group). (**G**) H&E staining of BAT, with quantification of lipid droplets area (µm²) (n = 5 per group). Scale bar = 100 µm. (**H**) Representative H&E staining of BAT, iWAT and eWAT. Scale bar = 50 µm. (**I**) TEM of BAT reveals enlarged mitochondria with disrupted cristae in *Tmem41b^Ucp1Cre^* mice. Scale bar = 1 µm. Right, Mitochondrial area quantification. (**J**) Measurement of basal, complex 1 (C1), complex 2 (C2) and maximal respiratory capacity (Maximum) in BAT from *Tmem41b^f/f^* and *Tmem41b^Ucp1Cre^* mice (n = 4 per group). (**K**) Western blot analysis of mitochondrial proteins SDHB and NDUFB8 in BAT (n = 3 per group). (**L**) RT-qPCR analysis of thermogenic genes in BAT (n = 4 per group). (**M**) Protein levels of UCP1 in BAT (n = 3 per group). (**N**) Schematic representation of the proposed role of *Tmem41b* in brown adipose tissue.

**Fig. S4.**
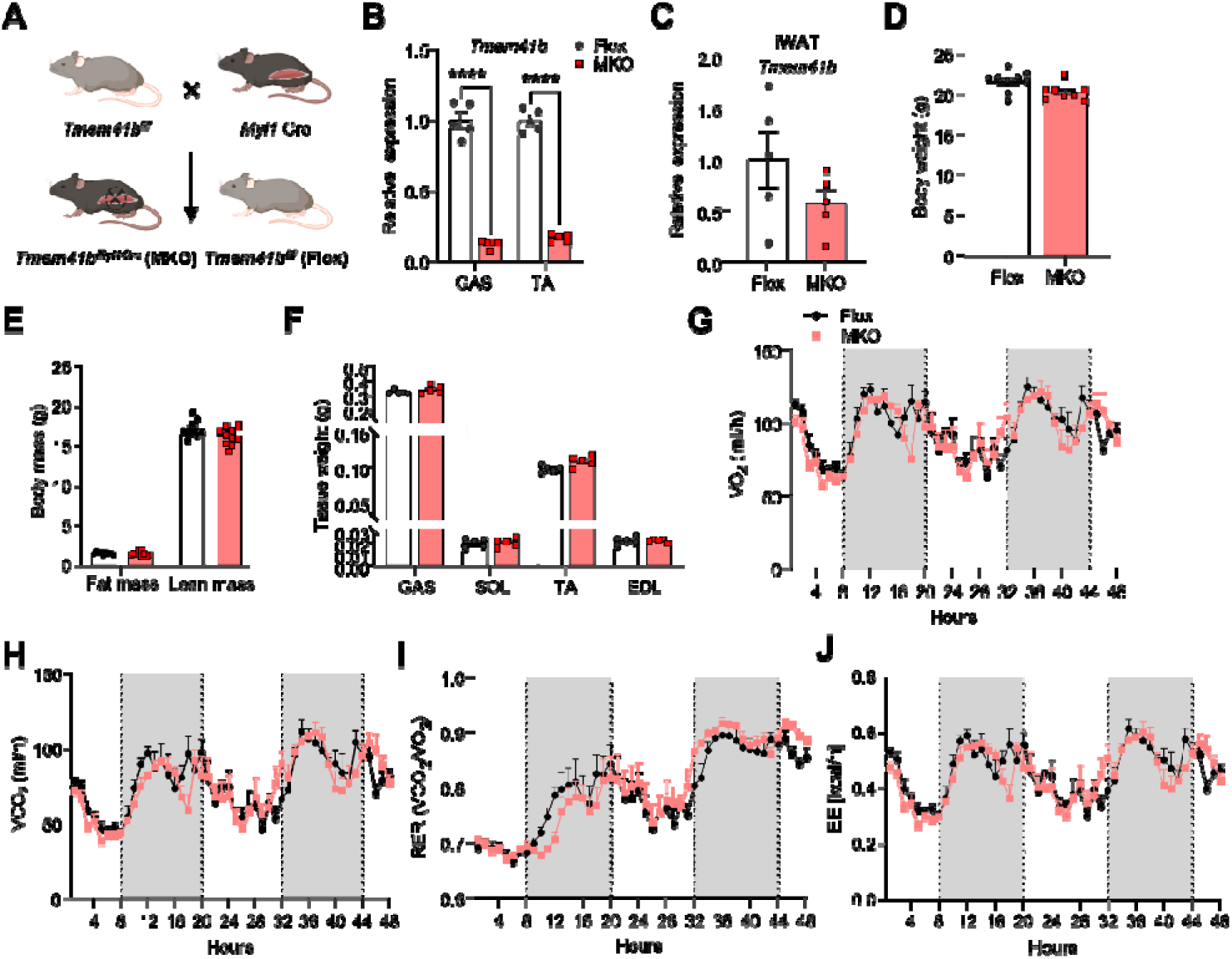
Mice with *Tmem41b* ablation in skeletal muscle have no effect on exergy expenditure. (**A**) Schematic diagram showing the strategy for generating skeletal muscle-specific *Tmem41b* knockout (*Tmem41b^Myl1Cre^*, MKO) mice. (**B**) Relative mRNA expression levels of *Tmem41b* in gastrocnemius (GAS) and tibialis anterior (TA) tissues from *Tmem41b^f/f^* and *Tmem41b^Myl1Cre^* mice, normalized to *tbp* (n = 5 per group). (**C**) Relative mRNA expression levels of *Tmem41b* in iWAT from *Tmem41b^f/f^* and *Tmem41b^Myl1Cre^*mice, normalized to *36b4* (n = 5 per group). (**D-E**) Body weight and body composition were assessed in 10-week-old *Tmem41b^f/f^* and *Tmem41b^Myl1Cre^* mice (n = 9, 8 per group). (**F**) Muscles weight of *Tmem41b^f/f^* and *Tmem41b^Myl1Cre^* mice (n = 5 per group). (**G-J**) Measurements of metabolic parameters of indicated mice. (G) Oxygen consumption (VO_2_); (H) Carbon dioxide production (VCO_2_); (I) Respiratory exchange ratio (RER); calculated as VCO_2_/VO_2_; (J) Energy expenditure (EE). Values represent mean ± SEM (n=5 mice per group).

**Fig. S5.**
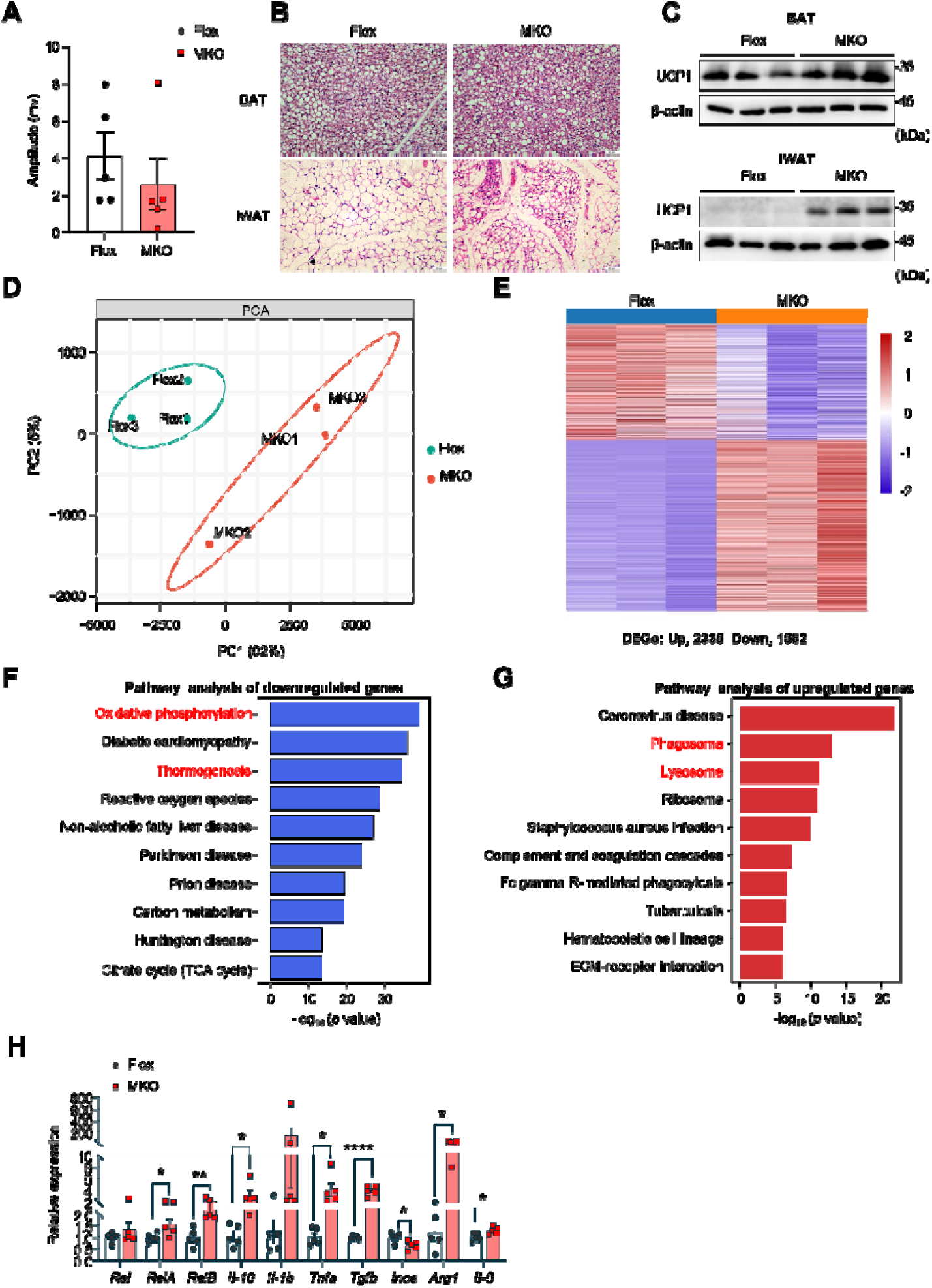
Mice with *Tmem41b* ablation in skeletal muscle are cold sensitive. (**A**) Amplitude was performed to evaluate muscle electrical activity in *Tmem41b^f/f^* and *Tmem41b^Myl1Cre^* mice (n = 5 per group). (**B**) Representative images of BAT and iWAT stained with H&E. Scale bar = 50 µm. (**C**) Western blots of UCP1 protein expression in BAT and iWAT (n = 3 per group). (**D**) PCA of RNA-seq data of TA muscles from *Tmem41b^f/f^* (Flox) and *Tmem41b^Myl1Cre^* (MKO) mice (n = 3 per group). (**E**) Heatmap of RNA-seq. (**F**) TOP 10 KEGG pathways enriched by downregulated genes. (**G**) TOP 10 KEGG pathways enriched by upregulated genes. (**H**) RT-qPCR analysis of the expression of inflammation-related genes (n = 5 per group).

**Fig. S6.**
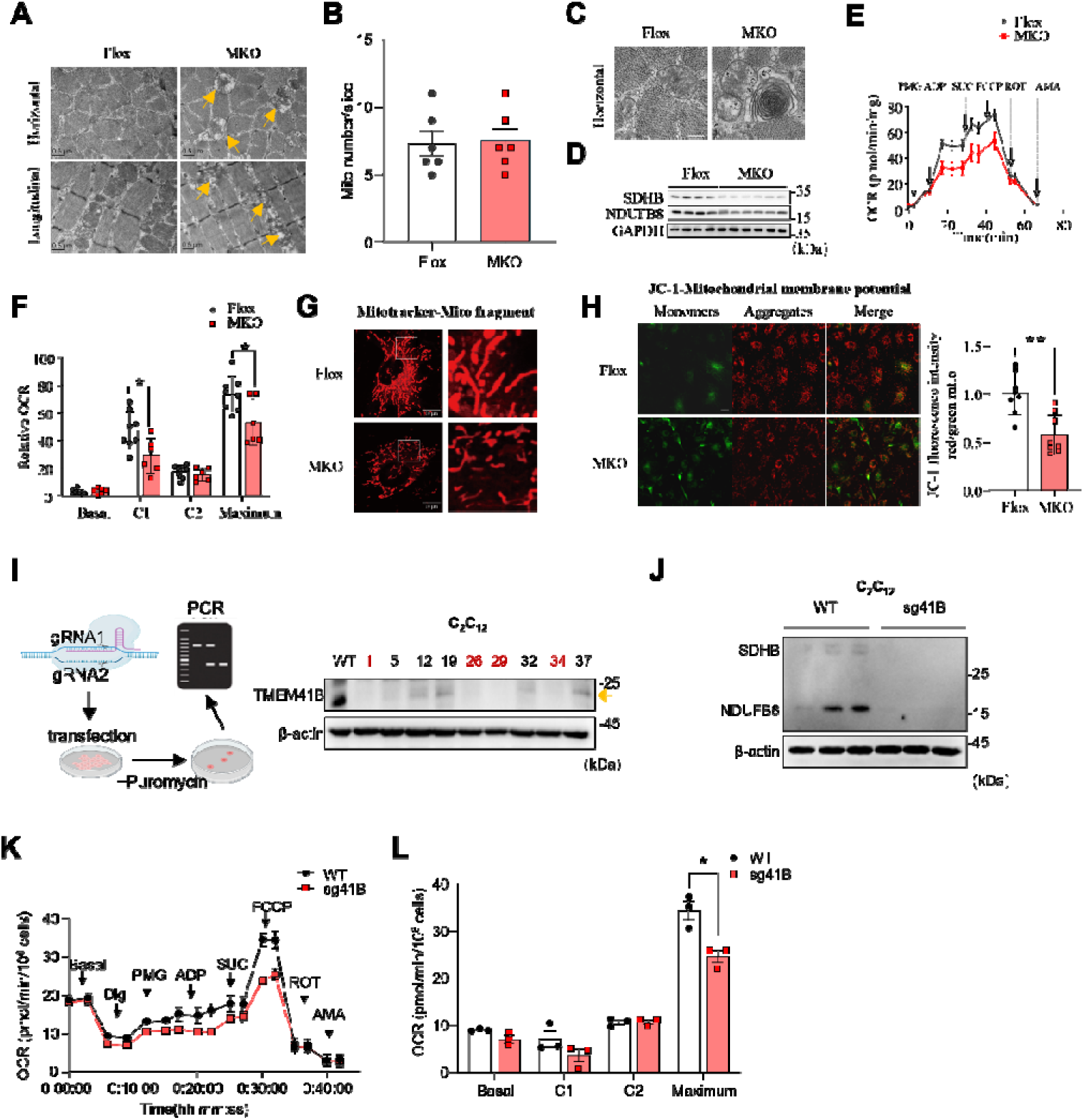
TMEM41B regulates mitochondrial functions in C_2_C_12_. (**A**) TEM of muscles in *Tmem41b^Myl1Cre^* mice. Yellow arrows, abnormal cristae. Scale bar = 0.5 µm. (**B**) The number of mitochondria per image was quantified (n = 6 per group). (**C**) Representative images of skeletal muscle. *Tmem41b^Myl1Cre^* mice showed an onion-like mitochondria. Scale bar = 0.5 µm. (**D**) Western blot analysis of mitochondria complexs SDHB and NDUFB8 in the indicated cells. (**E**) Representative traces of oxygen consumption rates (OCR) in C_2_C_12_ cells (n = 4 per group). (**F**) Quantification of basal, C1, C2 and Maximum (n = 3 per group). (**G**) Representative images of mitotracker staining in primary myocytes from *Tmem41b^Myl1Cre^*and control mice. Scale bar = 10 µm. (**H**) JC-1 staining in primary myocytes from *Tmem41b^f/f^* and *Tmem41b^Myl1Cre^* mice. Representative fluorescence images were shown, with quantification of the red-to-green fluorescence ratio as an indicator of mitochondrial membrane potential (n = 8 per group). Scale bar = 25 µm. (**I**) Generation and validation of TMEM41B knockout clonal cell lines. (**J**) Western blot analysis of mitochondrial proteins SDHB and NDUFB8 in C_2_C_12_ cells (n = 3 per group). (**K**) Representative traces of oxygen consumption rates (OCR) in C_2_C_12_ cells (n = 4 per group). (**L**) Quantification of basal, C1, C2 and Maximum (n = 3 per group).

**Fig. S7.**
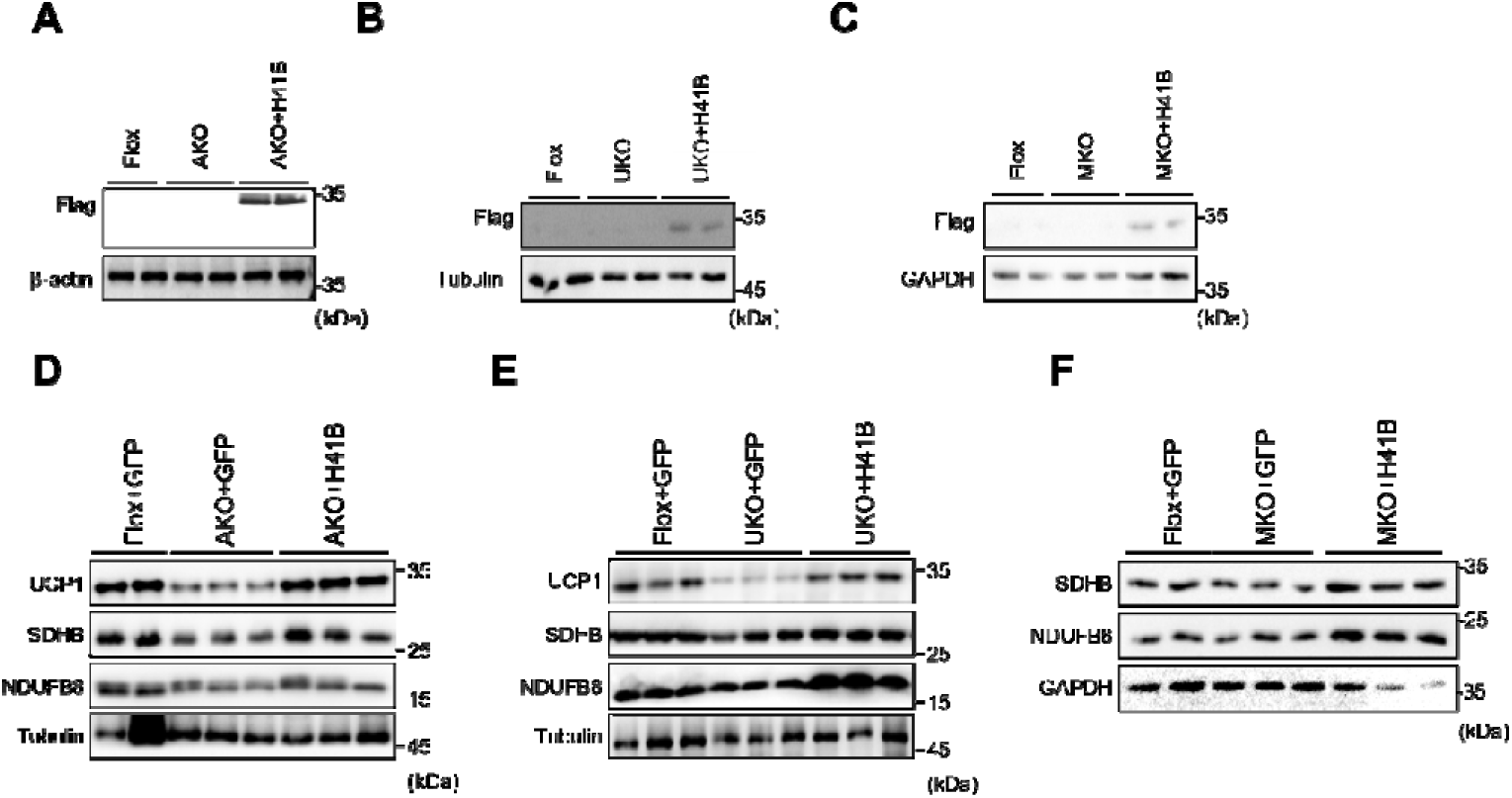
Tissue-specific reconstitution of human TMEM41B and its effects on mitochondrial complexes. **(A-C)** Western blot analysis of Flag-TMEM41B expression in BAT or muscle tissues bilaterally injected with AAV constructs encoding either puro vector or full-length human TMEM41B (H41B). (A) AKO mice model; (B) UKO mice model; (C) MKO mice model. (**D-F**) Representative immunoblots of mitochondrial complex proteins SDHB and NDUFB8 expression analysis in BAT tissue of AKO and UKO mice, or muscle of MKO mice (n=2-3 per group).

**Fig. S8.**
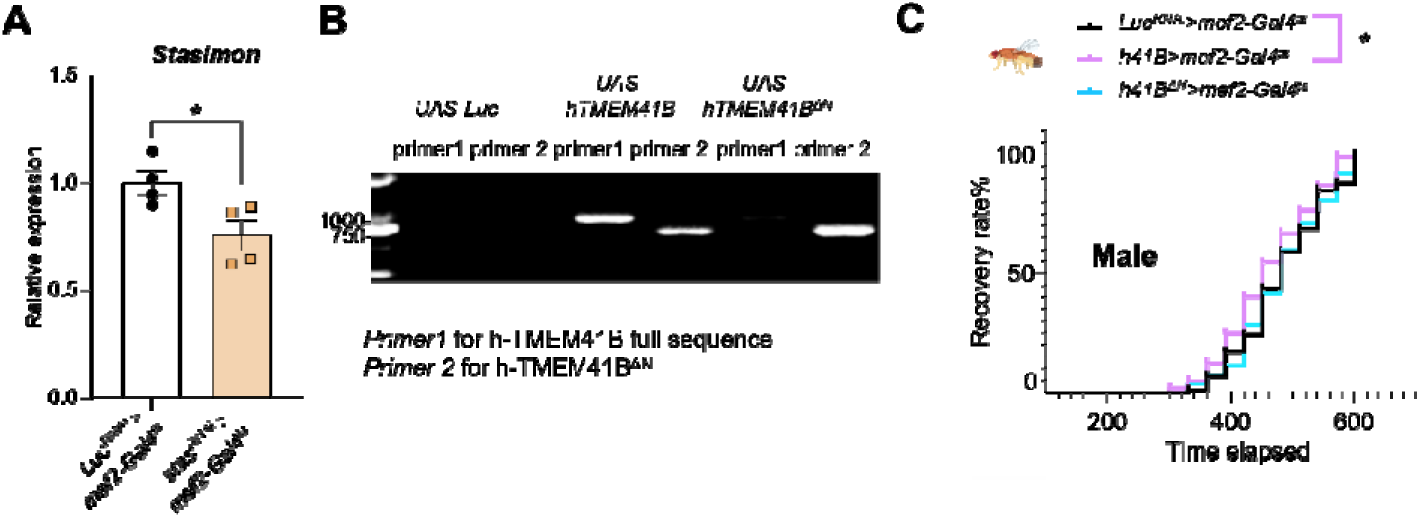
The N-terminal of human TMEM41B regulates cold tolerance in *drosophila*. (**A**) Relative mRNA expression levels of *stasimon* in muscles from *Luciferase^RNAi^* _>_ *mef2-Gal4^ts^* (*Luc^RNAi^* _>_ *mef2-Gal4^ts^*) and *Stasimon^RNAi^* _>_ *mef2-Gal4 ^ts^*(*Stas^RNAi^*_>_*mef2-Gal4^ts^*) *Drosophila*, normalized to *rp49* (n = 4 per group). (**B**) PCR confirmation of UAS-hTMEM41B and UAS-hTMEM*41B*^Δ*N*^ *Drosophila*. (**C**) Chill coma recovery for *Drosophila* (n > 100 per group).

**Fig. S9.**
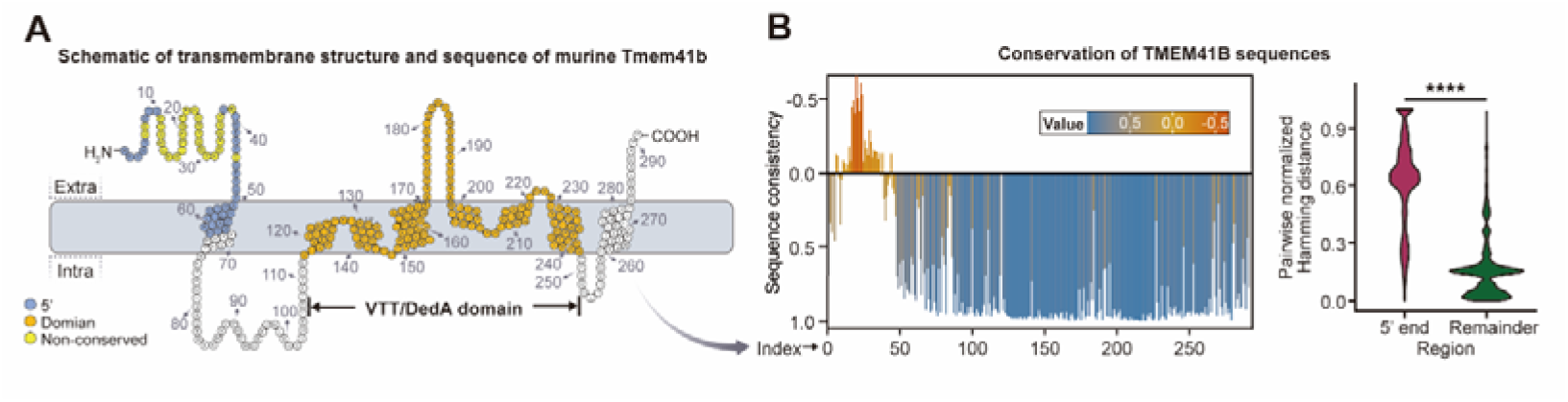
TMEM41B structure and its low-conservation N-terminal region. (A) Schematic diagram illustrating the transmembrane structure of murine *Tmem41b* in the endoplasmic reticulum. The N-terminal region (blue circles) is defined as the sequence preceding the potential new start codon at the first transmembrane loop (residue 68, M). Residues within the known VTT/DedA domain are marked with orange circles, and residues with a conservation score lower than 0 (panel B, left) are highlighted in yellow. The schematic is based on UniProt transmembrane structure prediction results and modified from the figure in Okawa et al (*33*). (B) TMEM41B N-terminus shows low cross-species conservation. (left) sequence consistency for each amino acid of TMEM41B across species. Lower consistency score indicates higher sequence variability. See Methods for the detailed consistency score definition. (right) Violin plot showing differential nucleotide conservations between the 5’-end region and the remaining sequences across vertebrates. The score is calculated using the Hamming distance between TMEM41B homologs from each pair of species. ****: *p* < 0.0001, determined by the paired t-test.

**Fig. S10.**
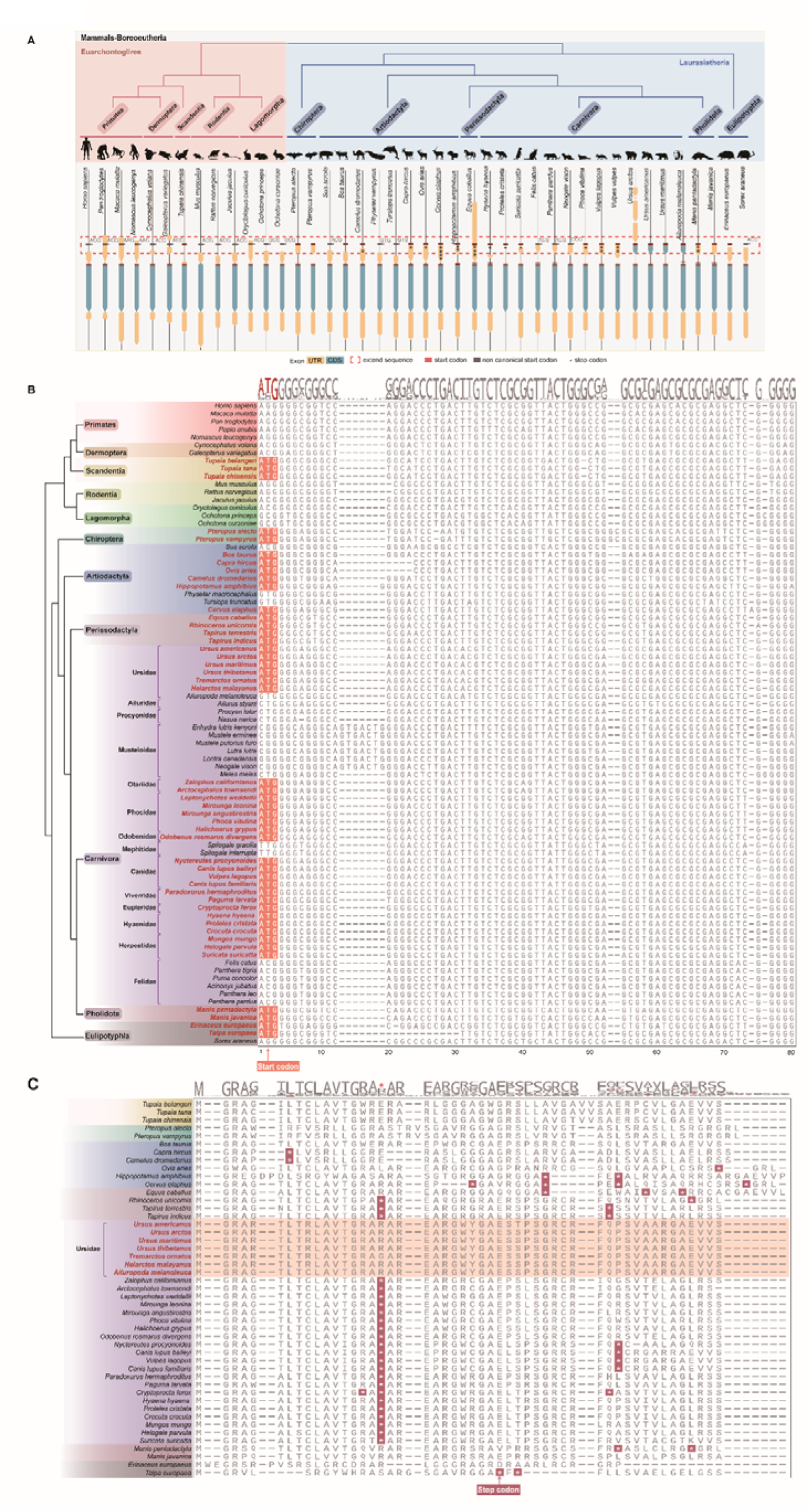
Evolutionary diversification of the TMEM41B N-terminal region across Boreoeutheria. (**A**) Extended version of Fig. 3A. Comparative gene structures of *Tmem41b* across the complete set of representative boreoeutherian species analyzed in this study. Each row corresponds to one species and illustrates the exon–UTR–CDS organization. A pronounced N-terminal coding-region extension is observed in *Ursidae* (red dashed box). Non-canonical start codons and premature stop codons are indicated by gray box and black asterisks (*), respectively. Complete species lists and genomic sequence sources are provided in table S3. (**B**) Multiple sequence alignment of the first 80 nucleotides of the *Tmem41b* 5′ region from 82 representative mammalian species. The alignment highlights the canonical start codon (ATG) region and identifies the species in which this start codon is retained. (**C**) Amino acid sequence alignment of species that retain a canonical ATG start codon (encoding methionine). The alignment shows the N-terminal translated region, with lineage-specific amino acid substitutions and premature stop codons highlighted.

**Fig. S11.**
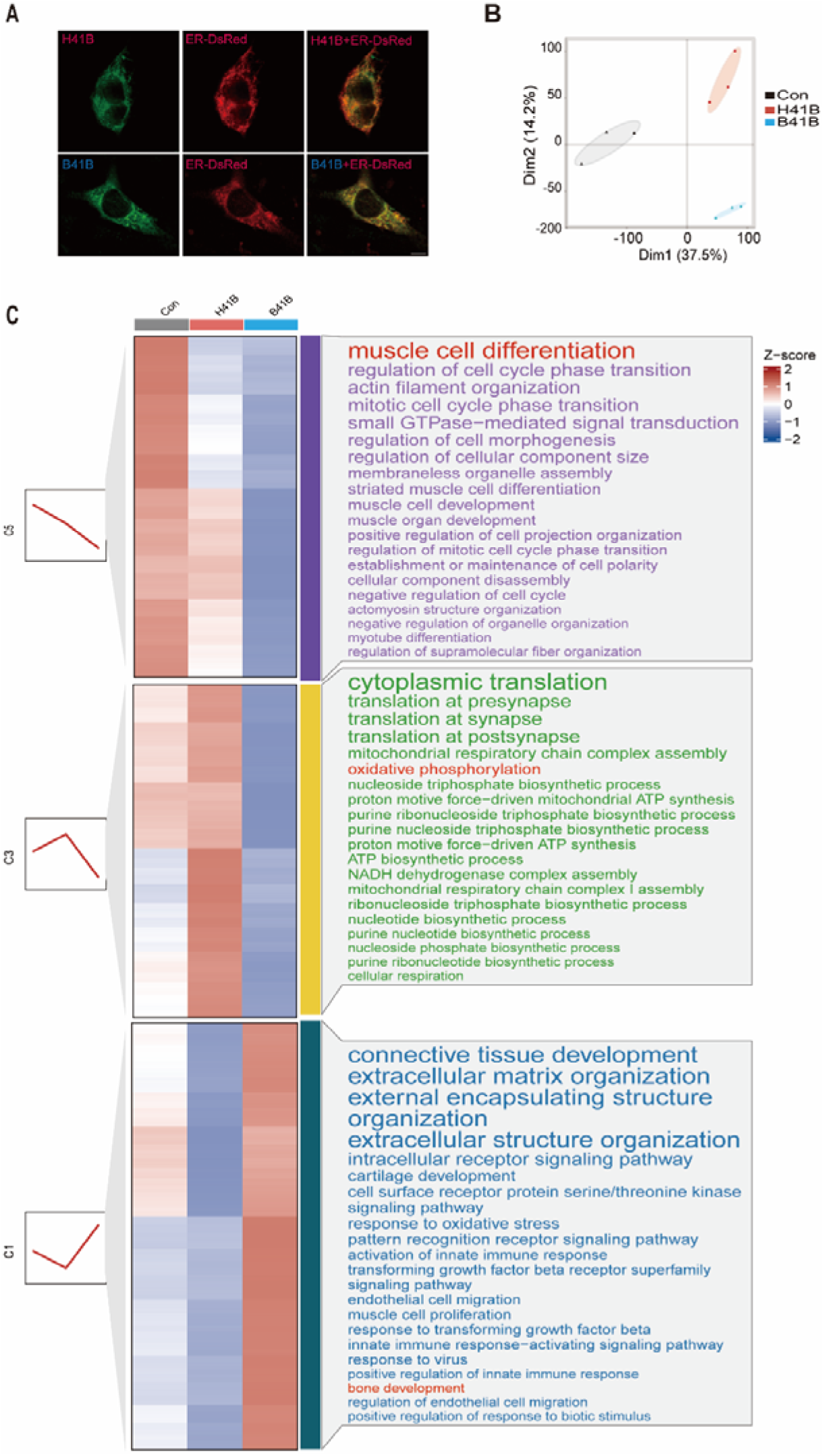
Overexpression of human TMEM41B or bear TMEM41B in C_2_C_12_ cells. (**A**) Representative subcellular localization microscopy images of both H41B and B41B colocalizes with ER. Scale bar = 10 µm. (**B**) PCA of RNA-seq data from overexpression of H41B and B41B in C_2_C_12_ cells and control cells (n = 3 per group). (**C**) Heatmap of RNA-seq data in three distinct patterns of change with showing top 20 GO pathways for each cluster.

**Fig. S12.**
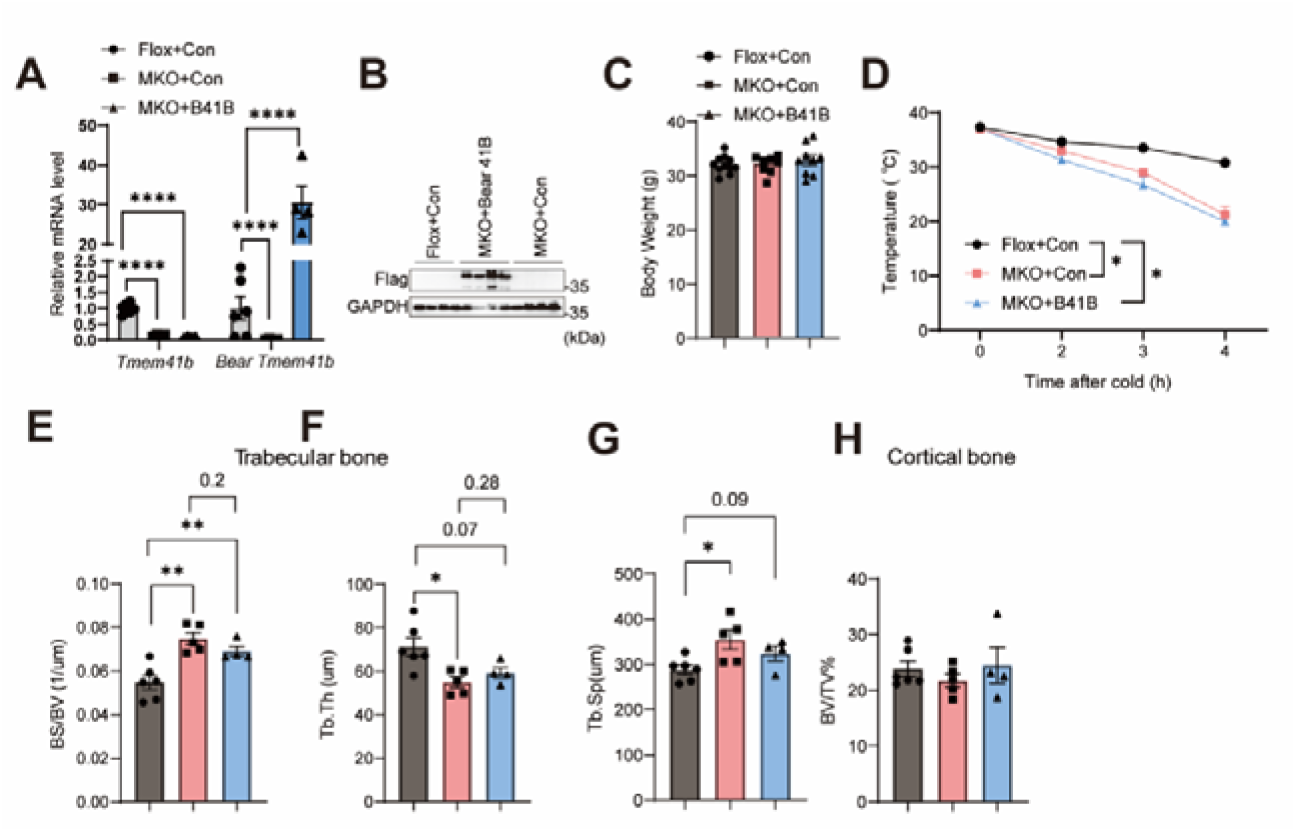
51-amino-acids substitution induces a torpor-like state in mice. **(A)** Relative mRNA expression levels of N terminal *Tmem41b* and *Tmem41b* in muscles of from *Tmem41b^f/f^*, *Tmem41b^Myl1Cre^* and *Tmem41b^Myl1Cre+B41B^* mice, normalized to tbp (n = 5 per group). (**B**) Representative immunoblot confirming ectopic expression of Flag-TMEM41B protein in the skeletal muscle of the indicated mice (n = 3 per group). (**C**) Body weight analysis of mice models (n = 8 to 11 per group). (**D**) Core body temperature of mice from indicated groups during a 4-hour cold challenge (4L) (n = 9 per group). (**E-F**) Quantification of trabecular bone micro-architecture. (E) Bone Surface / Bone Volume (BS/BV); (F) Trabecular thickness (Tb.Th). (**G-H**) Analysis of cortical bone micro-CT. (**G**) Trabecular Separation (Tb.sp); (**H**) Percentage of Bone Volume/Total Volume (BV/TV%).

