## Supplementary figures and images for "A Bear-Specific Coding Exon in TMEM41B Is Associated with Cold Adaptation"

### Supplemental figure

#### Slide 1
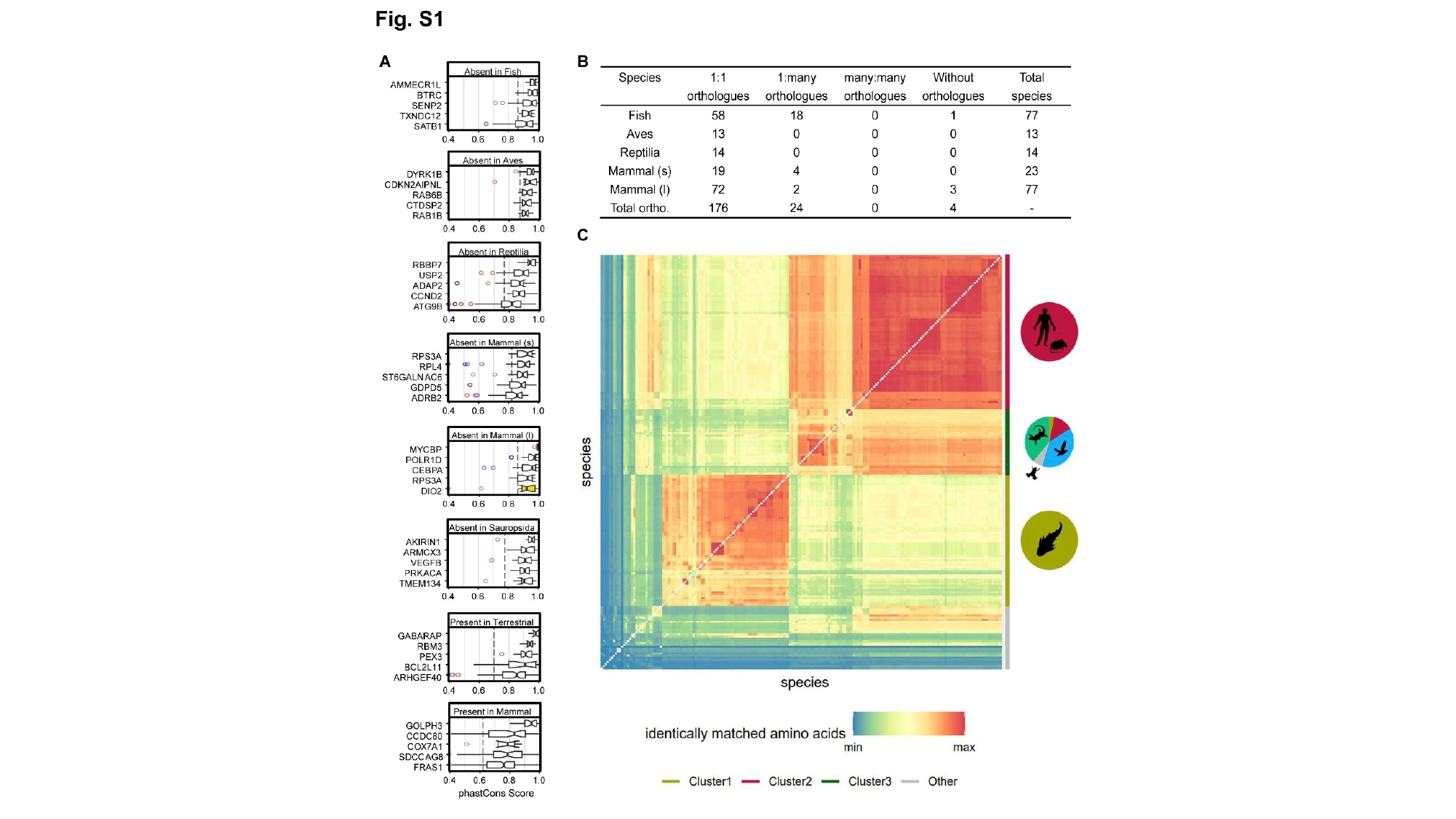

#### Slide 2
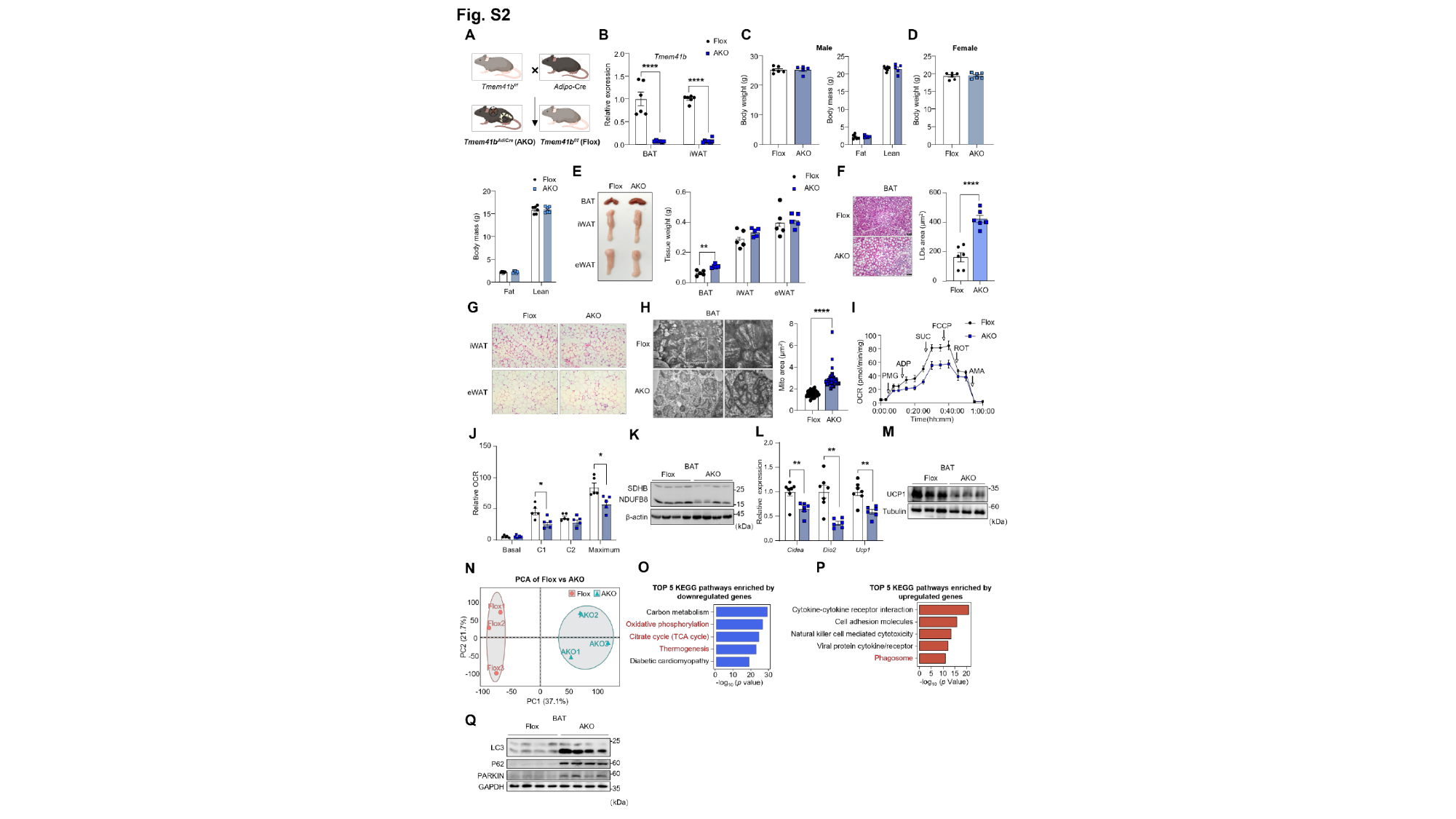

#### Slide 3
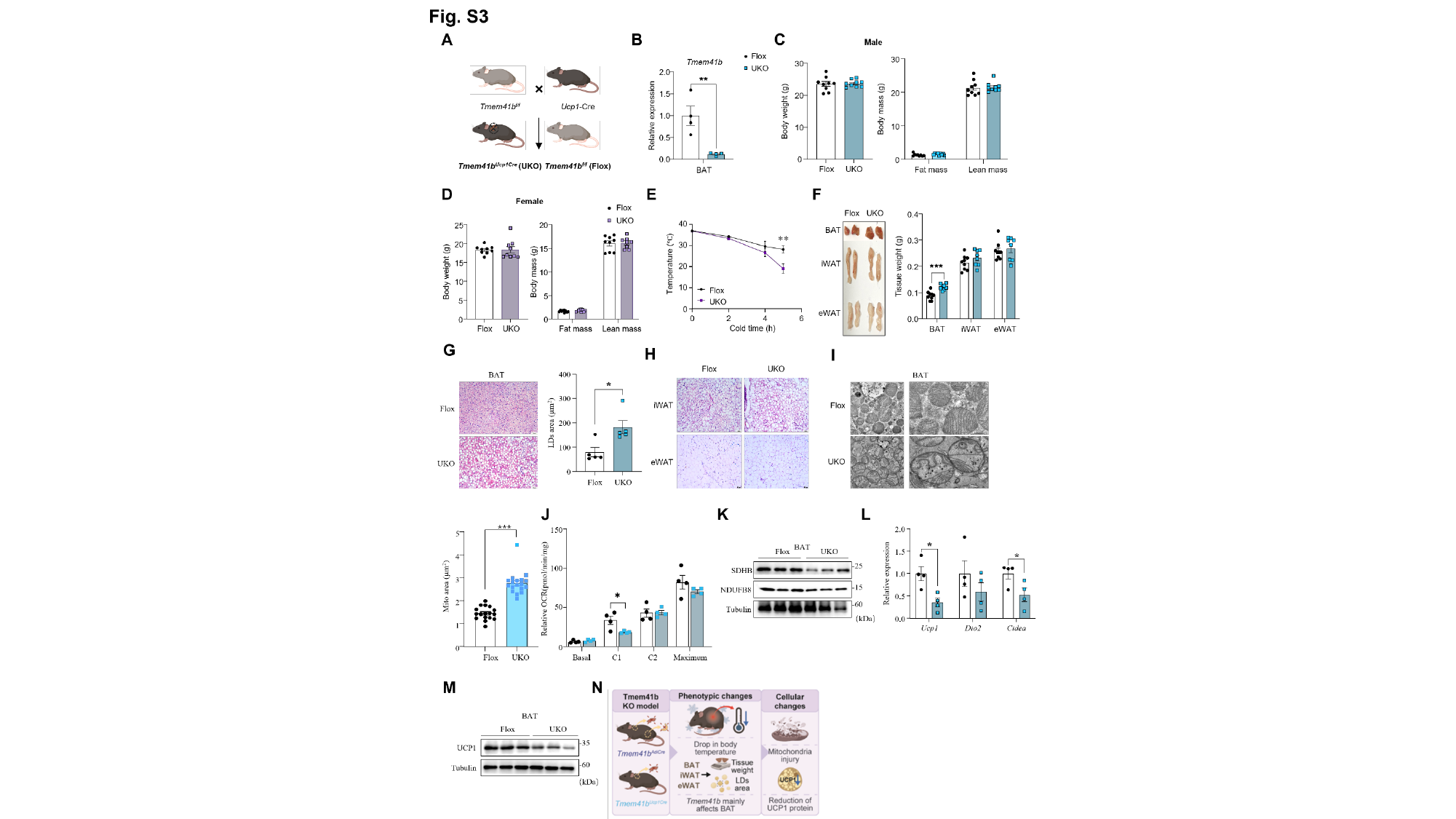

#### Slide 4
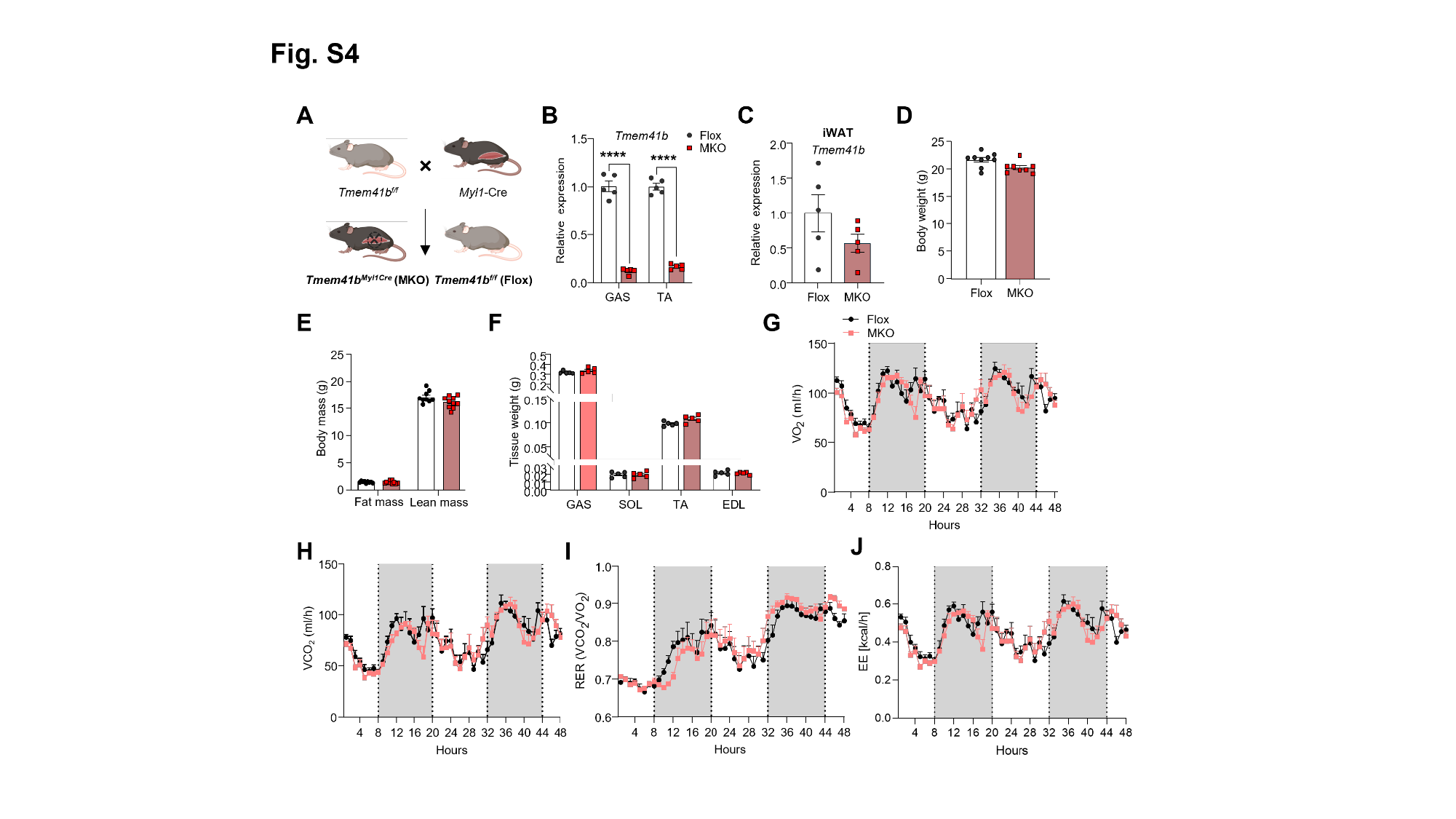

#### Slide 5
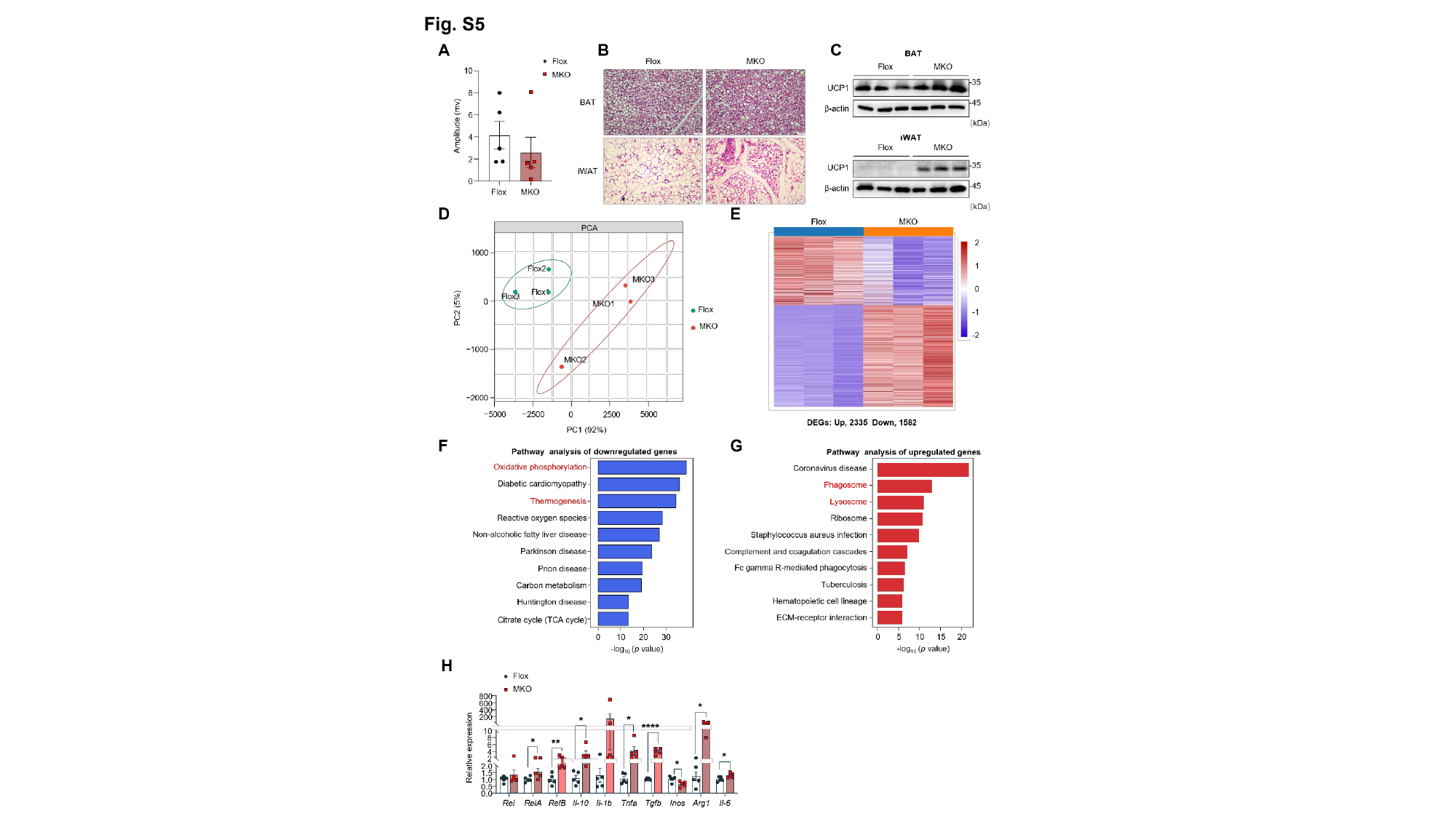

#### Slide 6
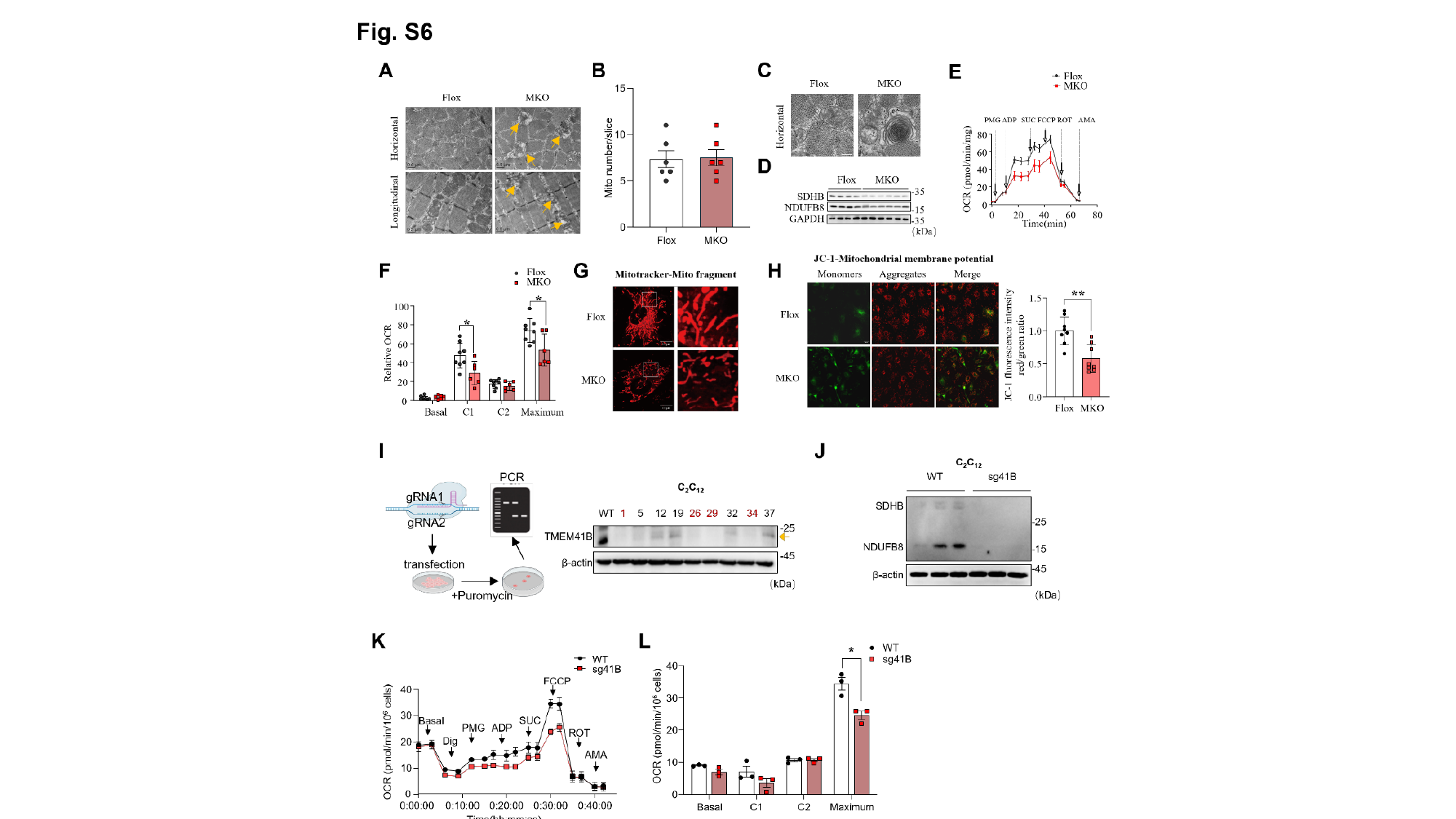

#### Slide 7
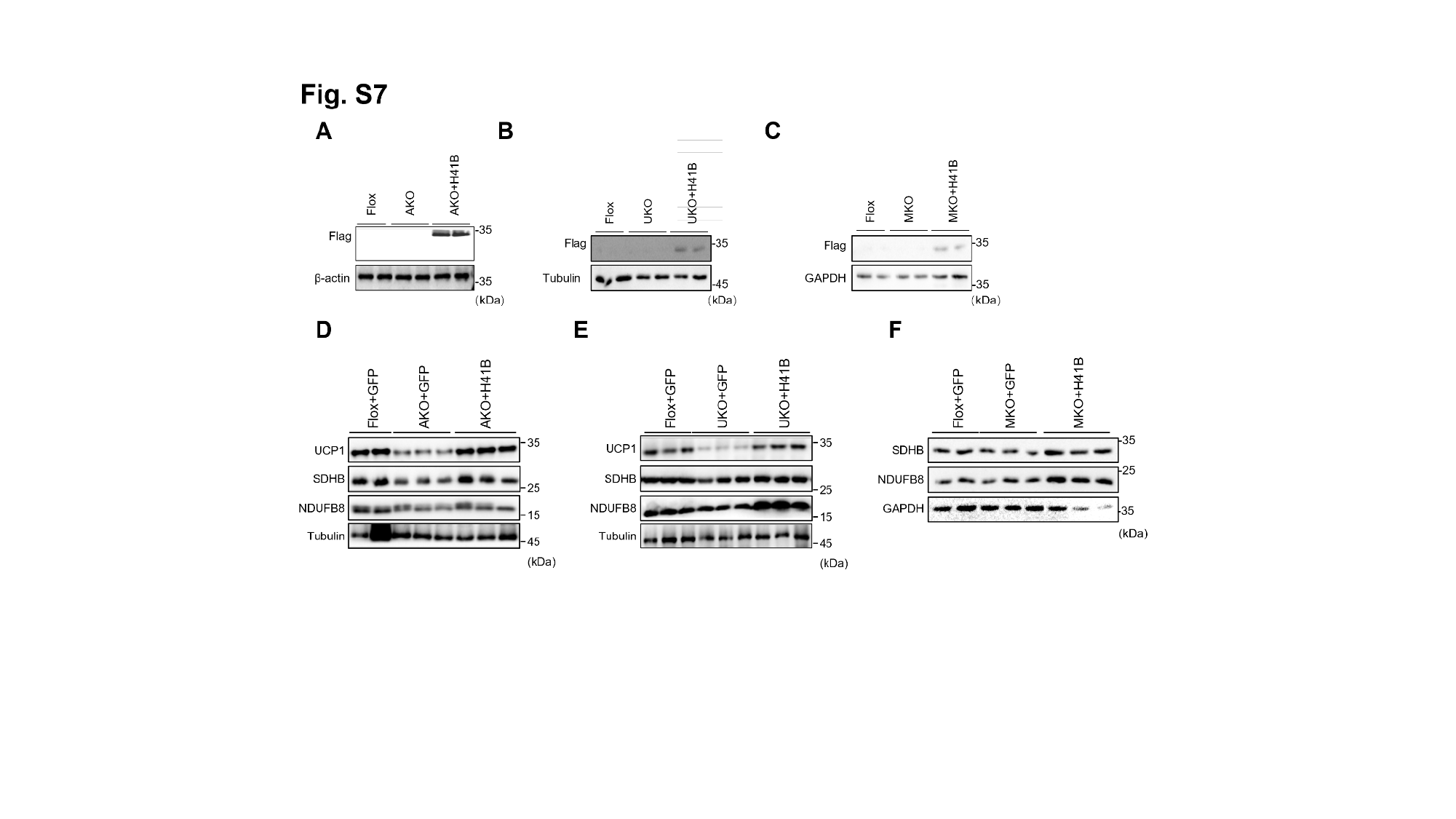

#### Slide 8
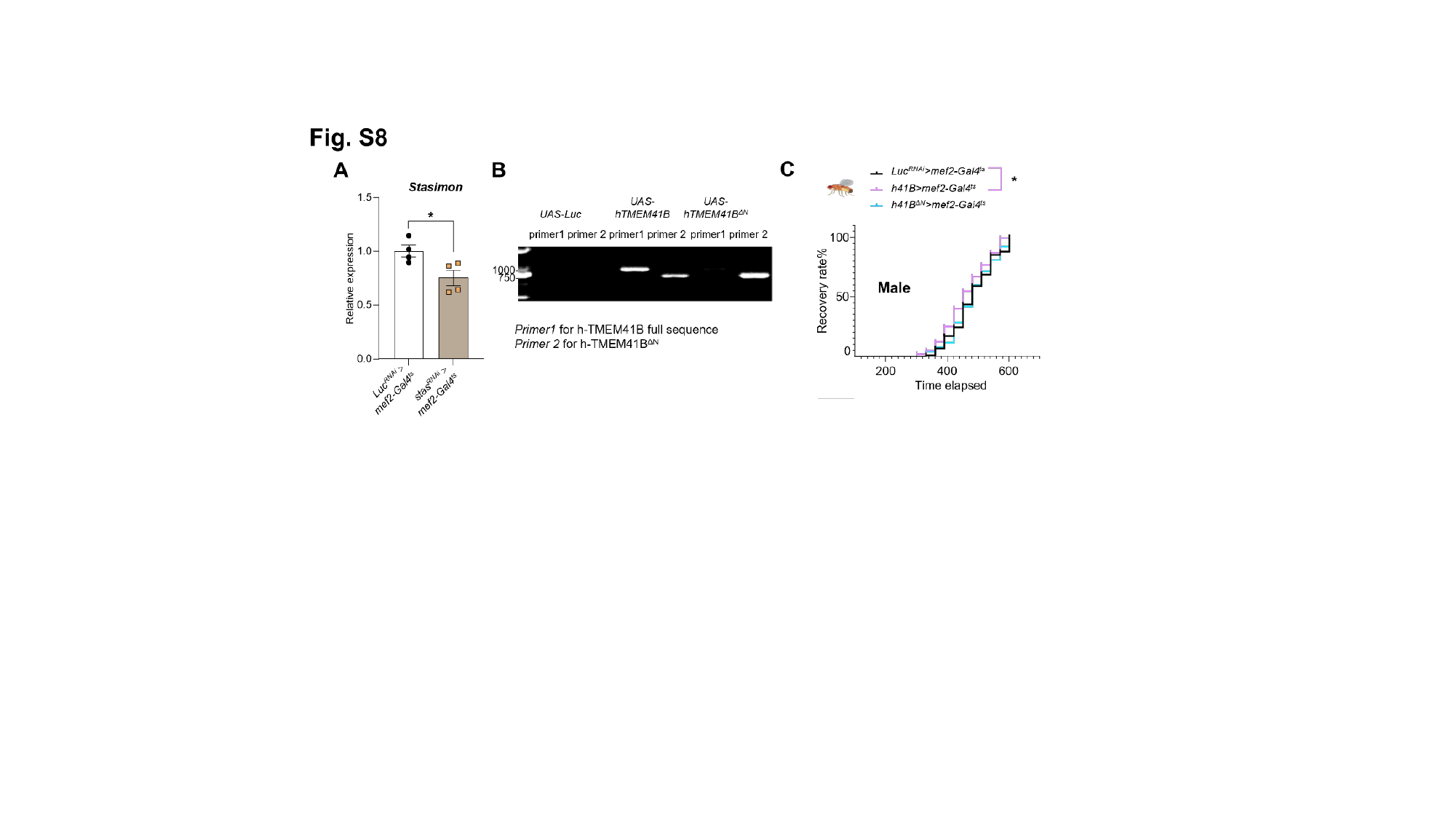

#### Slide 9
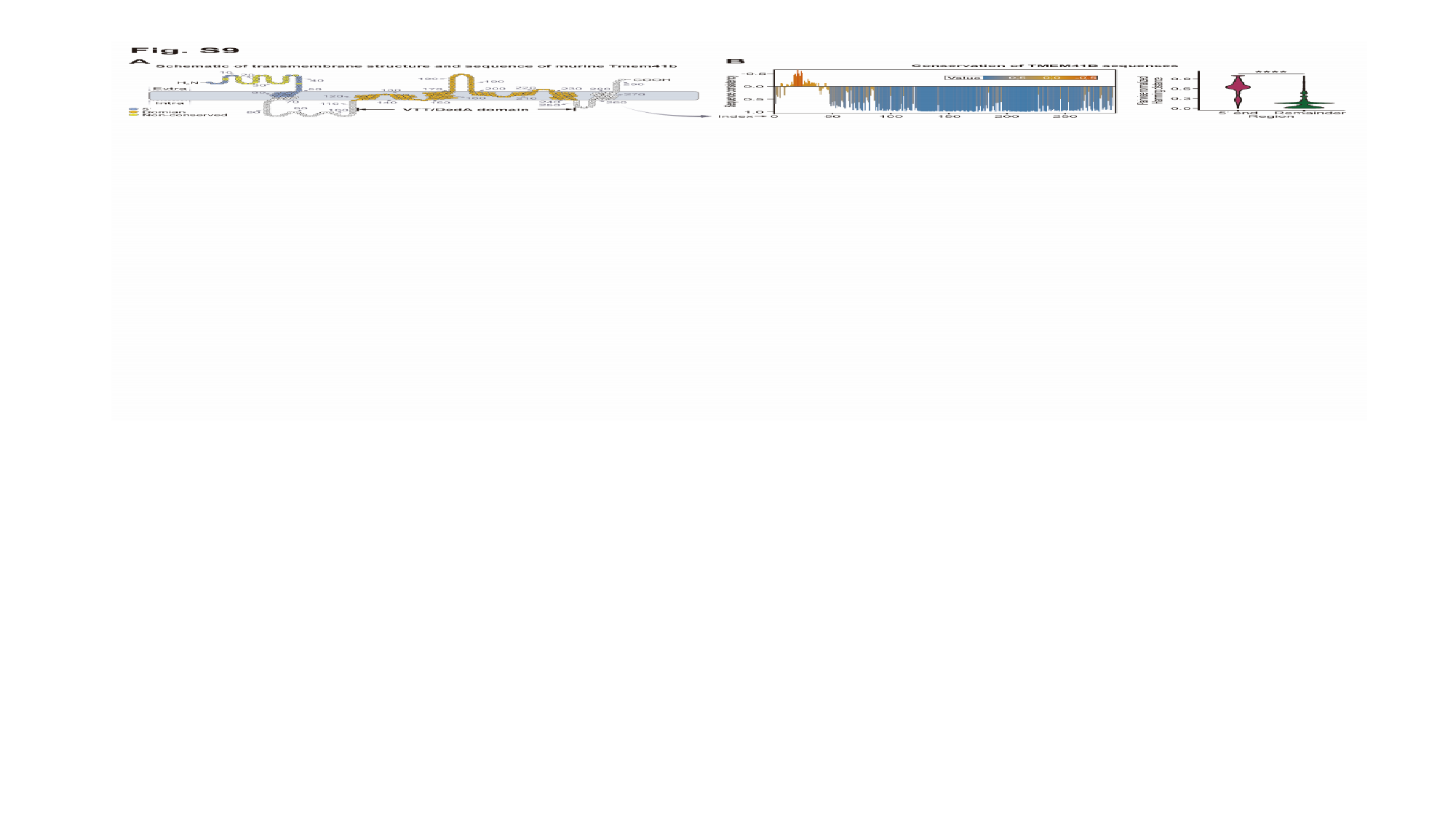

#### Slide 10
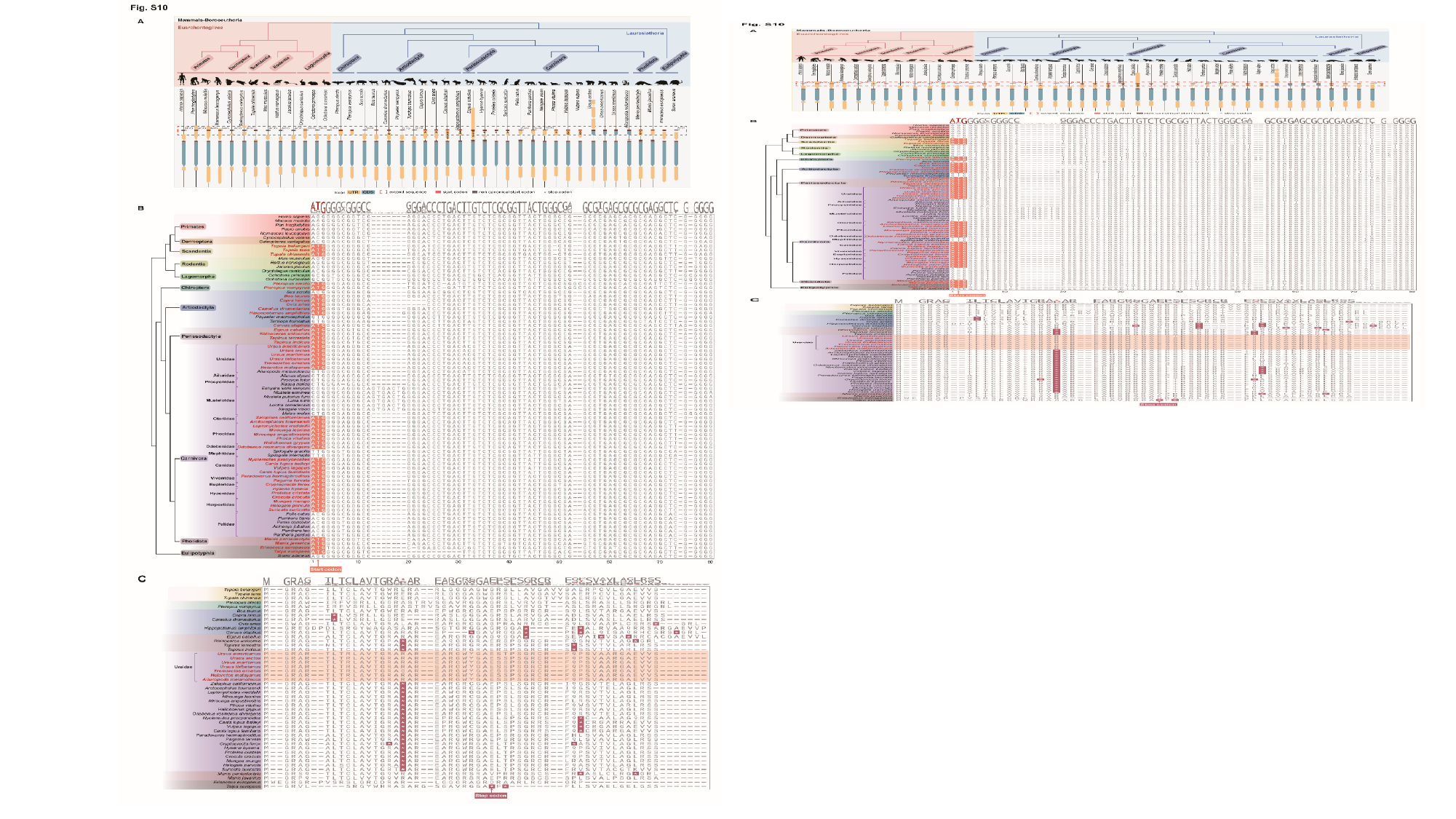

#### Slide 11
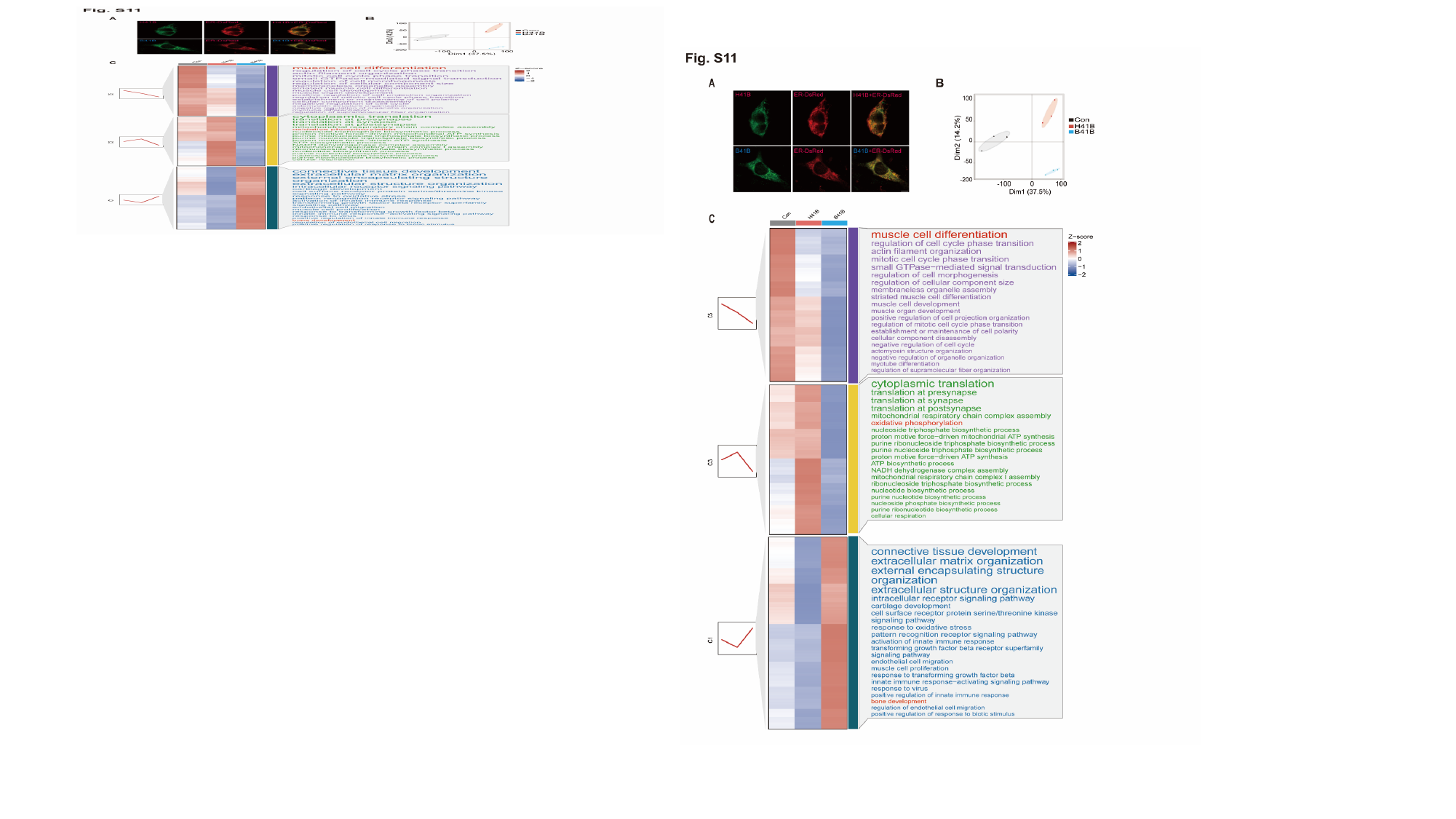

#### Slide 12
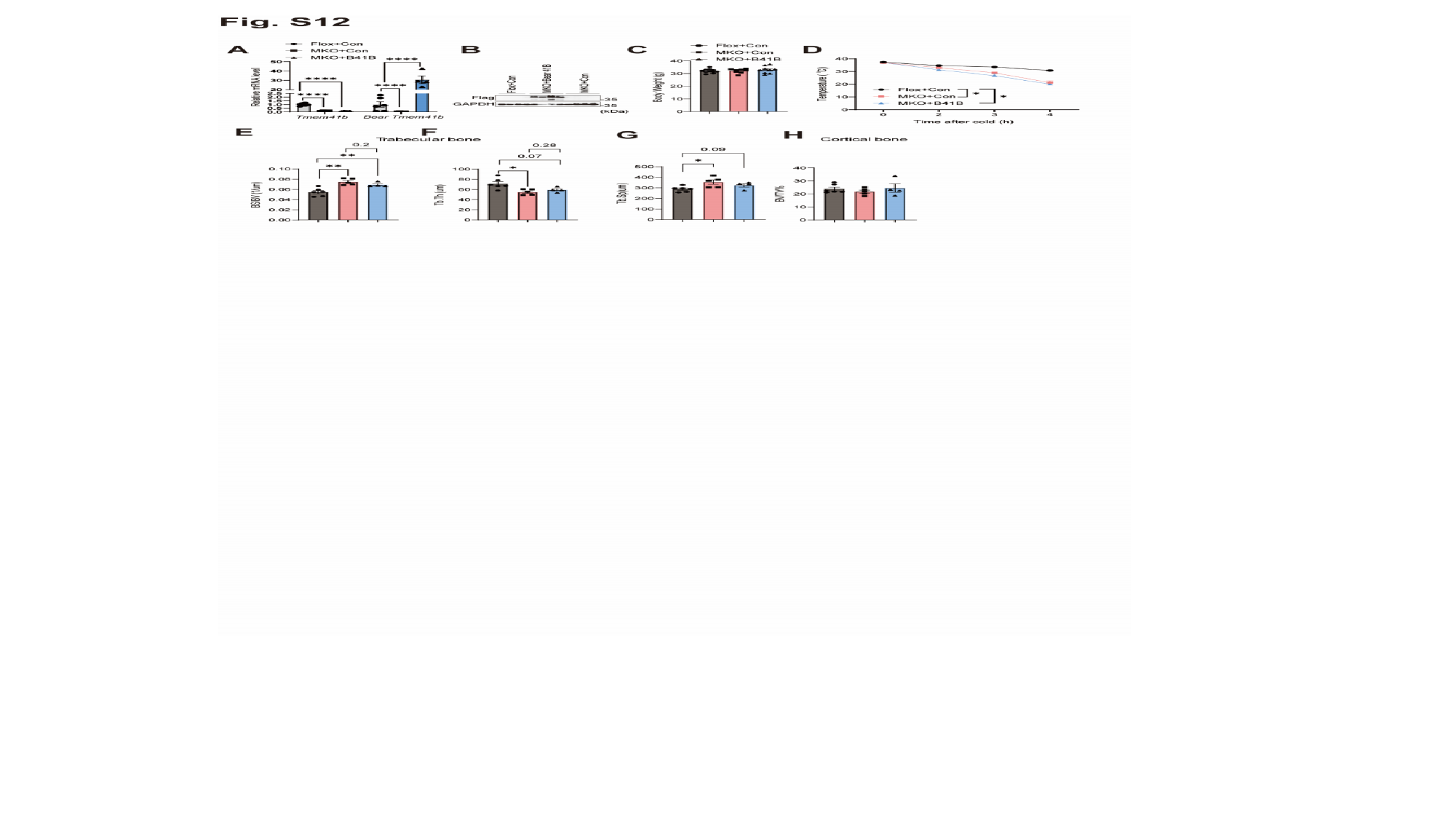
